# Pleiotropic Meta-Analysis of Cognition, Education, and Schizophrenia Differentiates Roles of Early Neurodevelopmental and Adult Synaptic Pathways

**DOI:** 10.1101/519967

**Authors:** Max Lam, W. David Hill, Joey W. Trampush, Jin Yu, Emma Knowles, Gail Davies, Eli Stahl, Laura Huckins, David C. Liewald, Srdjan Djurovic, Ingrid Melle, Kjetil Sundet, Andrea Christoforou, Ivar Reinvang, Pamela DeRosse, Astri J. Lundervold, Vidar M. Steen, Thomas Espeseth, Katri Räikkönen, Elisabeth Widen, Aarno Palotie, Johan G. Eriksson, Ina Giegling, Bettina Konte, Annette M. Hartmann, Panos Roussos, Stella Giakoumaki, Katherine E. Burdick, Antony Payton, William Ollier, Ornit Chiba-Falek, Deborah K. Attix, Anna C. Need, Elizabeth T. Cirulli, Aristotle N. Voineskos, Nikos C. Stefanis, Dimitrios Avramopoulos, Alex Hatzimanolis, Dan E. Arking, Nikolaos Smyrnis, Robert M. Bilder, Nelson A. Freimer, Tyrone D. Cannon, Edythe London, Russell A. Poldrack, Fred W. Sabb, Eliza Congdon, Emily Drabant Conley, Matthew A. Scult, Dwight Dickinson, Richard E. Straub, Gary Donohoe, Derek Morris, Aiden Corvin, Michael Gill, Ahmad R. Hariri, Daniel R. Weinberger, Neil Pendleton, Panos Bitsios, Dan Rujescu, Jari Lahti, Stephanie Le Hellard, Matthew C. Keller, Ole A. Andreassen, Ian J. Deary, David C. Glahn, Anil K. Malhotra, Todd Lencz

**Author notes:** Lead Contact and Correspondence: Todd Lencz, Zucker Hillside Hospital, Division of Psychiatry Research, 75-59 263rd Street, Glen Oaks, NY, 11004, USA.

## Abstract

Liability to schizophrenia is inversely correlated with general cognitive ability at both the phenotypic and genetic level. Paradoxically, a modest but consistent positive genetic correlation has been reported between schizophrenia and educational attainment, despite the strong positive genetic correlation between cognitive ability and educational attainment. Here we leverage published GWAS in cognitive ability, education, and schizophrenia to parse biological mechanisms underlying these results. Association analysis based on subsets (ASSET), a pleiotropic meta-analytic technique, allowed jointly associated loci to be identified and characterized. Specifically, we identified subsets of variants associated in the expected (“Concordant”) direction across all three phenotypes (i.e., greater risk for schizophrenia, lower cognitive ability, and lower educational attainment); these were contrasted with variants demonstrating the counterintuitive (“Discordant”) relationship between education and schizophrenia (i.e., greater risk for schizophrenia and higher educational attainment). ASSET analysis revealed 235 independent loci associated with cognitive ability, education and/or schizophrenia at p<5×10^−8^. Pleiotropic analysis successfully identified more than 100 loci that were not significant in the input GWASs, and many of these have been validated by larger, more recent single-phenotype GWAS. Leveraging the joint genetic correlations of cognitive ability, education, and schizophrenia, we were able to dissociate two distinct biological mechanisms: early neurodevelopmental pathways that characterize concordant allelic variation, and adulthood synaptic pruning pathways that were linked to the paradoxical positive genetic association between education and schizophrenia. Further, genetic correlation analyses revealed that these mechanisms contribute not only to the etiopathogenesis of schizophrenia, but also to the broader biological dimensions that are implicated in both general health outcomes and psychiatric illness.

## Introduction

It has long been observed that impaired cognitive ability is a significant aspect of the illness in schizophrenia^1–5^. Cognitive deficits have been shown to be largely independent of clinical state and treatment status in patients with schizophrenia^1,4,6–9^, and are observed (in more subtle form) in their first-degree relatives^10,11^. Moreover, cognitive deficits precede illness onset by many years, beginning in early childhood^5,12–14^, resulting in reduced educational attainment in patients diagnosed with schizophrenia^15,16^. Recent advances in psychiatric and cognitive genomics have reliably demonstrated that the inverse relationship between cognitive ability and risk for schizophrenia is also observed at the molecular genetic level (r_g_∼-.20) ^17–23^. Paradoxically, genetic correlation studies have indicated a *positive* relationship between educational attainment and risk for schizophrenia (r_g_∼.10)^20,22,24–27^, despite the fact that educational attainment and cognitive ability exhibit a very strong polygenic overlap (r_g_∼.70)^19,20,27^. Educational attainment is often considered to be a proxy for cognitive ability; however, the lack of perfect genetic overlap between the two, combined with the paradoxical genetic correlation between educational attainment and schizophrenia, suggest an opportunity to decompose distinct genetic mechanisms accounting for this pattern of results.

Whereas genetic correlation analysis has recently become widespread due to the availability of techniques such as linkage disequilibrium (LD) score regression (LDSC)^25,28^, these approaches generally result in a single, genome-wide estimate of polygenic overlap. Moreover, novel meta-analytic approaches (e.g., Multi-Trait Analysis of GWAS [MTAG]^29^) for merging seemingly heterogeneous GWAS datasets tend to exploit commonalities across phenotypes rather than differences; for example, two recent studies have employed MTAG across the highly correlated cognitive and educational GWASs in order to accelerate the process of gene discovery^19,20^. By contrast, few studies have attempted to examine the counter-intuitive correlation between schizophrenia and educational attainment, or to parse subsets of SNPs that might drive cross-phenotype correlations. An initial effort has successfully identified a few individual loci that act in paradoxical fashion, increasing educational attainment while simultaneously increasing risk for schizophrenia^30^; two other studies have identified loci that demonstrate other pleiotropic effects^21,31^.

To date, however, no studies have utilized pleiotropic meta-analytic techniques to comprehensively parse variance from cognitive, educational, and schizophrenia GWAS that may pinpoint differential biological mechanisms. In order for the paradoxical pattern of genomewide correlations to exist, there must be identifiable subsets of SNPs that are differentially involved in driving these genetic relationships. Therefore, we sought to identify differentially associated variants, as they may yield crucial insights into the fine-grained genetic architecture of schizophrenia, in turn giving us insights into the etiopathogenic mechanisms underlying the illness that standard GWAS cannot detect.

In the present report, we first utilize a simple subsetting approach to identify SNPs that are significantly associated with *either* cognitive ability or educational attainment, but not both (Supplementary Figure 1a). We hypothesized that these SNP subsets would demonstrate stronger genetic correlations with schizophrenia than observed with a simple genomewide approach. We then employ a pleiotropic meta-analytic approach, ASSET^32^, which permits the characterization of each SNP with respect to its pattern of effects on multiple phenotypes (Supplementary Figure 1b). For example, ASSET has previously been used to demonstrate that the minor allele of rs2736100 (at the *TERT* locus) is positively associated with risk for pancreatic cancer, negatively associated with risk for kidney and lung cancers, and not significantly associated with risk for cancers of the breast, bladder, and prostate; other cancer loci were demonstrated to have various other patterns of effects^32^. We utilized ASSET to identify two types of loci: 1) those SNPs that are consistently associated with all three phenotypes in the expected direction (i.e., the same allele associated with higher cognitive ability, higher educational attainment, and lower risk for schizophrenia), which we label “Concordant”; and 2) SNPs that demonstrate the paradoxical association between education and schizophrenia (i.e., the same allele associated with higher educational attainment and higher risk for schizophrenia), labelled “Discordant.” Next, we compared the statistically significant ASSET results to the output of single trait GWASs of cognitive ability, educational attainment, or schizophrenia^22,35,36^, in order to identify novel loci suggested by ASSET. Subsequently, we conducted a series of pathway and transcriptome-wide analyses to biologically characterize differential mechanisms underlying Concordant vs Discordant loci. Finally, we performed a series of genetic correlation analyses to compare the overlap of Concordant and Discordant SNP subsets with other relevant traits. Further analytic details are covered in the methodology section; the full analysis workflow is also represented in Supplementary Figures 1-2.

## Methods

### Stage 1: Simple Subsetting Approach Based on P-values in Cognition and Education GWAS

Note that for most purposes in this manuscript, we are using the largest GWAS for schizophrenia^33^, cognitive ability^19^, and educational attainment^34^ published prior to 2018. For each of these phenotypes, larger GWAS have been published in 2018; these were used for validation and extension as described in subsequent sections. Before any subsetting analyses, genomewide genetic correlations using LD score regression^25,37^ were used to confirm the earlier observed genetic correlations between schizophrenia with both cognitive ability and education. In Stage 1, preliminary SNP subsets were formed simply based on p-value thresholds of cognitive ability and educational attainment GWAS: i) SNPs nominally associated with cognition (*p* < 0.05) and not associated with education (*p* > 0.05) were selected, resulting in 74,470 SNPs; ii) SNPs nominally associated with cognition (*p* < 0.05) and not associated with education using a stricter threshold (p > 0.5) resulted in 66,657 SNPs; iii) similar procedures were carried out for SNPs nominally associated with education (*p* < 0.05) but not cognition (*p* > 0.05), resulting in 104,807 SNPs; and iv) SNPs nominally associated with education (*p* < 0.05) and not cognition using a stricter threshold (*p* > 0.5), resulting in 44,803 SNPs.

Next, we performed a heterogeneity test between results of cognitive and educational GWAS using METAL^38^, and generated sets of SNPs showing opposite effects between the two (i.e, the same allele predicts better cognitive performance but less educational attainment, and vice versa). We identified sets of SNPs of varying sizes based on varying p-value thresholds for the heterogeneity test (p < .5; p < .25; p < .1; p < .05; p < .01; p < .001).

To evaluate the degree of genetic correlation of these preliminary subsets of SNPs with respect to schizophrenia, we utilized GNOVA^39^, a recently published method similar to LD score regression. However, GNOVA is specifically designed for examination of genetic correlations using SNP subsets (rather than full genomewide summary statistics), whereas such applications have not been explicitly tested in LD score regression and may not be robust.

### Stage 2: ASSET Meta-Analysis and ASSET-generated SNP Subsets

Schizophrenia GWAS summary statistics were obtained from the Psychiatric Genomics Consortium^33^ based on the European ancestry GWAS of schizophrenia (N = 77,096, Cases = 33,640, Controls = 43,456; GWAS mean χ^2^ = 1.677). To make them compatible with effect sizes (Beta weights) derived from the linear regression-based cognition and education GWASs, odds-ratios from the case-control schizophrenia GWAS were converted to Beta by taking the natural logarithm. Effect direction per SNP was also reversed for schizophrenia to make them consistent with the interpretation of cognition and education (i.e., concordant alleles are those where the direction of effect is the same for cognitive ability, educational attainment, and for schizophrenia). Summary statistics for the education GWAS was obtained from the Social Science Genomics Association Consortium^34^ (N = 328,917, GWAS mean χ^2^ =1.638). GWAS summary statistics for cognition are available via earlier inverse-variance meta-analysis of samples^19^ from the COGENT^27^ consortium (N = 107,207, GWAS mean χ^2^ = 1.245). General quality control parameters were applied to the schizophrenia and cognitive GWAS summary statistics excluding SNPs with INFO scores < 0.6 and minor allele frequency < 0.01; multiple quality control parameters thresholds were reported for the education GWAS^34^ and summary statistics were used as provided on http://www.thessgac.org/data Detailed quality control and meta-analytic procedures were reported previously^19^. Only SNPs that were present for all three phenotypes were retained as input to the ASSET meta-analysis, resulting in 7,306,098 SNPs for subsequent analysis.

Pooling GWAS summary statistics using conventional inverse-variance meta-analysis increases power, but also poses methodological challenges when different studies are capturing heterogeneous/pleiotropic phenotypes. In the case of pleiotropy, individual variants are likely to be associated with only a subset of the traits analyzed, or may even demonstrate effects in different directions for the different phenotypes under analysis. ASSET meta-analysis^32^ is an agnostic approach that generalizes standard fixed-effects meta-analysis by allowing a subset of the input GWASs to have no effect on a given SNP, and exhaustively searches across all possible subsets of “non-null” GWAS inputs within a fixed-effect framework to identify the strongest association signal in both positive and negative directions. ASSET then evaluates the significance of these positive and negative associations while accounting for multiple testing. This methodology allows for a powerful pooled two-tailed Z-score test statistic that effectively combines p-values for variants with strong effects in opposite directions across input GWASs. ASSET also permits the addition of a covariance term for the adjustment of overlapping samples. More recently, comparisons between cross phenotype meta-analysis methodologies demonstrated that ASSET performed best as the number of meta-analysed traits with null effects increases, along with specificity and sensitivity of the results; in addition the ASSET approach best controlled for potential Type 1 inflation due to sample overlap, and non-uniform distribution of effect sizes^40^. As such we selected ASSET for its conservative effect estimates and minimal inflation for the purpose of the current report.

GWAS summary statistics from schizophrenia, cognition, and education were combined using ASSET two-tailed meta-analysis (version 1.9.1) to obtain a single cross phenotype pleiotropic GWAS results. Default parameters were carried out using the “h.traits” function. Inter-study correlations of the phenotype were first ascertained using linkage disequilibrium score regression (LDSC)^25,28^, which accounts for the genetic correlation of the phenotypes as well as sample overlap. For each given SNP, ASSET generates Z-scores of effect size and p-values based on the strongest association from the input studies in positive and negative directions, respectively; then these p-values are pooled into a single two-tailed p-value for pleiotropy^32,40,41^. SNPs with similar relationships across the input traits (regardless of statistical significance) are then grouped into subsets identified by ASSET (see Figure 1b; Supplementary Figure 1, bottom).

Again, as noted above, it is important to emphasize that the per-SNP direction of effect was reversed for schizophrenia to make it consistent with the interpretation of cognition and education (i.e., higher scores are better, such that higher scores for schizophrenia are now coded as *decreased* risk for the disorder). Thus, in the notations to follow, a ‘∩’, represents variant subsets with the same e?ect directions, following the reversal of the direction of effect for the schizophrenia data set, and ‘|’,represent traits that goes in the opposite direction in terms of effect sizes compared to the other two traits (again following the reversal of the direction of effect for the schizophrenia data set). ASSET subsets included: i) scz ∩ edu ∩ cog (Concordant, variants with an allele associated with an increase in cognitive ability and educational attainment, but a decrease in schizophrenia risk); ii) edu ∩ cog | scz (Schizophrenia outlier, variants associated with an increase in cognitive ability and educational attainment, but also an increase in schizophrenia risk); iii) scz ∩ cog | edu (Education outlier, variants associated with an increase in schizophrenia risk and reduced cognitive ability, but an increase in educational attainment); iv) scz ∩ edu | cog (Cognition outlier, variants associated with an increase in schizophrenia risk and reduced educational attainment, but an increase in cognitive ability) subsets. ASSET also identified SNPs where only a single trait (scz or edu or cog) was significant; these were included in a category called ‘Single phenotype’.

Finally, to generate an appropriate contrast for the ‘Concordant’ subset, we included a combined single ‘Discordant’ subset, representing the counter-intuitive genetic correlation between education and schizophrenia, where, ‘Discordant’ = (edu ∩ cog | scz) + (scz ∩ cog | edu) [subsets ii and iii above], regardless of the effect for cognition. These contrasts are also represented visually in Figure 1 and Supplementary Figure 1.

### Consolidation of Independent Loci

Independent genomewide significant loci for each ASSET meta-analysis subset were identified via SNP clumping procedures that are a part of the Functional Mapping and Annotation (FUMA) pipeline^42^. For the LD-rich MHC region, a single top SNP was retained. For all other loci, clumping procedures were carried out based on the European 1000 Genomes Project phase 3 reference panel. First, “independent significant SNPs” were defined as those SNPs with a p-value < 5×10^−8^ and with linkage disequilibrium r^2^ < 0.6 with other, more significant SNPs at the same locus. Second, candidate SNPs were then identified for subsequent annotations and were defined as all SNPs that had a MAF of 0.01, a maximum p value of 0.05, and were in LD of ≥ r^2^ 0.6 with at least one of the independent significant SNPs. To ensure that biological annotation of these loci would not be hampered by poor coverage at any locus, candidate SNPs included SNPs from the 1000 Genomes reference panel that may not have been included in the ASSET data. Third, “lead SNPs” were defined as the independent significant SNPs that had the strongest signal at a given locus and were strictly independent from each other (r^2^ < 0.1). Finally, risk loci that were 250kb or closer were merged into a single locus. The FUMA procedure was iterated across all ASSET SNP subsets, which were comprised (by definition) of non-overlapping SNPs. Additional variant annotations were conducted with ANNOVAR^43^, and lookups with published GWAS were conducted using the GWAS catalog^44^. Additional SNP lookups were performed with the input summary statistics (Cognition, Education and Schizophrenia GWASs^33,34,19^), recent MTAG analysis of Intelligence^20^, recent Cognition/Intelligence^22,23^ GWASs, and pleiotropic analyses of cognition and schizophrenia^21^ as well as education and schizophrenia^31^. RAggr^45^ was utilized to extract SNPs within 250kb and r^2^ > 0.6 from published reports to allow merging from loci generated from ASSET subsets.

### MAGMA Gene-based Analysis: Tissue Expression and Competitive Pathway Analysis

SNPs were mapped to 19,436 protein coding genes using MAGMA, as implemented in the FUMA^42^ pipeline. MAGMA^46^ gene analysis was performed using default SNP-wide mean model using the 1000 Genomes phase 3 reference panel and default gene annotations that are part of the FUMA pipeline. Genome-wide SNP p-values and SNP-level sample sizes were included in the input files. MAGMA gene tissue expression analysis was carried out utilizing the Genotype-Tissue Expression (GTEx; version 7^47–49^) resource, examining the relationship between gene expression in a specific tissue type and ASSET results. The gene-property test was performed for average expression (log2 transformed RPKM with pseudocount 1 after winsorization at 50) of 53 specific tissue types conditioning on average expression across all tissue types.

MAGMA competitive pathway analysis was also conducted with results that emerged from the ASSET analysis in order to identify specific biological processes linked to our sub-phenotypes of interest. Gene sets that were tested included drug-related pathways^50,51^, as well as custom-curated neurodevelopmental and other brain-related gene sets that had gone through stringent quality control in a study originally designed to interrogate rare variants in schizophrenia^52^. In the latter, pathways with more than 100 genes from Gene Ontology (release 146; June 22, 2015 release), KEGG (July 1, 2011 release), PANTHER (May 18, 2015 release), REACTOME (March 23, 2015 release), DECIPHER Developmental Disorder Genotype-Phenotype (DDG2P) database (April 13, 2015 release) and the Molecular Signatures Database (MSigDB) hallmark processes (version 4, March 26, 2015 release) were initially included. Additional gene sets were selected based on risk for schizophrenia and neurodevelopmental disorders, including those reported for schizophrenia rare variants^53^ (translational targets of FMRP^54,55^, components of the post-synaptic density^56,57^, ion channel proteins^57^, components of the ARC, mGluR5, and NMDAR complexes^57^, proteins at cortical inhibitory synapses^58,59^, targets of mir-137^57^, and genes near schizophrenia common risk loci^57,60^) and autism risk (autism risk genes: targets of *CHD8*^61–63^, splice targets of RBFOX^63–65^, hippocampal gene expression networks^66^, neuronal gene lists from the Gene2cognition database [http://www.genes2cognition.org]^63^, as well as “loss of function intolerant” genes (pLI > 0.9 from the ExAC v0.3.1 pLI metric), ASD risk genes for FDR < 10% and 30%, and ASD/developmental disorder *de novo* genes hit by a LoF or a LoF/missense *de novo* variant^67,68^). Brain-level tissue expression gene-sets included the Brainspan RNA-seq dataset^69^ and the GTEx v7 dataset^47^.MAGMA gene-based and gene-set analysis were conducted using MAGMA v1.6^46^.

### S-PrediXcan: Brain Tissue Expression Profiles and Hypergeometric Gene-set Enrichment Analysis

Genetically regulated gene expression was imputed for the ASSET summary statistics using tissue models from GTEx v7 and the CommonMind Consortium via S-PrediXcan (formerly MetaXcan)^70–72^. S-PrediXcan computes downstream phenotypic associations of genetic regulation of molecular traits using elastic nets, adjusting for model uncertainty and colocalization of GWAS and eQTL signals^73^. GTEx v7 tissue included amygdala (N = 88), anterior cingulate cortex (N=109), basal ganglia (N=144), cerebellar hemisphere (N=125), cerebellum, cortex (N=136), frontal cortex (N=118), hippocampus (N=111), hypothalamus (N=108), nucleus accumbens (N=130), putamen (N=111), spinal cervical-1 (N=83) and substantia nigra (N=80). The CommonMind consortium data consist of tissue expression derived from the dorsolateral prefrontal cortex (DLPFC, N=279)^74^. GTEX v7 Tissue expression models were trained using elastic net models that were made publicly available here (http://predictdb.org/). Elastic net models for DLPFC were contributed by collaborators from the CommonMind Consortium^74–76^. Bonferroni correction was first conducted for each ASSET subset of genes. Genes that survived multiple testing correction were entered to GENE2FUNC to conduct a test of over-representation, which is part of the FUMA^42^ pipeline. This analysis differs from the MAGMA gene set analysis as the MAGMA gene set analysis is used to examine if gene sets, united by a known biological theme, are enriched for the phenotype under investigation. In a test of overrepresentation, as conducted using GENE2FUNC, the shared function of the genes of interest is unknown, and its elucidation is the goal of a test of over-representation. The over-representation test conducted using GENE2FUNC queries gene sets from the 1) Molecular Signature Database (MsigDB v 5.2) 2) WikiPathways (Curated Version 20161010) and 3) GWAS Catalogue (reported genes ver e91 20180206); to avoid spurious results, we required a minimum of 3 genes per pathway. For each gene set, hypergeometric tests were conducted, examining the list of genes significant in each ASSET subset for overlap with gene sets within the databases stipulated above and applying Bonferroni correction for multiple testing. To reduce the likelihood that hypergeometric pathway analysis would be influenced by the dense number of genes within the MHC region, genes within the coordinates of 28,000,000 – 35,000,000^77^ on chromosome 6 were removed.

### Genetic Correlations

To examine how our ASSET Concordant and Discordant SNP subsets relate to other phenotypes with available GWAS data, we conducted genetic correlations using GNOVA^39^, an approach similar to LD score regression but capable of working with SNP subsets. A series of neuropsychiatric, inflammatory, brain, metabolic and cardiovascular phenotypes that have been previously demonstrated to have genetic correlations with cognitive measures were selected to interrogate the genetic overlaps of our ASSET subsets. Interpretation of GNOVA for the Concordant subset was straightforward, as the three input GWAS weights all follow the same direction (following the reverse-coding of schizophrenia as noted previously). By contrast, Discordant SNPs have two separate potential weights (allelic β for schizophrenia vs. allelic β for education); as shown in Figure 1B, a given SNP might have somewhat different effect sizes (distance from the center line) for education as compared to schizophrenia.

Therefore, we weighted each SNP by the stronger value of β: variants for which the schizophrenia β was stronger than the education β were referred to as “schizophrenia type,” while variants with the opposite pattern were referred to as “education type.” Nevertheless, it is important to emphasize that the Discordant SNPs represent a single dimension of biology, and the net effects of all “schizophrenia type” variants were equivalent to the “education type” SNPs, albeit with opposite signs.

## Results

### Stage 1: Preliminary evaluation of genetic correlations

GWAS summary statistics for cognition (N = 107,207)^19^ and education (N = 328,917)^34^ were used to evaluate preliminary genetic correlations with schizophrenia (N = 77, 096)^33^. Consistent with previous results, the inverse genetic correlation between cognition and schizophrenia was significant (r_g_ = -.19, se = .03, p=2.85×10^−10^), as was the counter-intuitive positive correlation between education and schizophrenia (r_g_=.10, se=.02, p=3.91×10^−5^). Note that these analyses were conducted prior to reversing the direction of effect for schizophrenia.

Prior to the main ASSET analysis, two simple approaches were used to examine subsets of SNPs and their association with schizophrenia (Supplementary Figure 1). First, we selected SNPs that were nominally associated with education (*p* < 0.05) and generally not associated with cognition (*p* > 0.05); GNOVA for this subset of educational attainment SNPs revealed a slightly stronger positive correlation r_g_ of .15 with schizophrenia (Figure 2a). With a stricter threshold for SNPs not associated with cognition (*p* > 0.50), these “non-cognitive” educational attainment SNPs attained an r_g_ of .20 with schizophrenia. GNOVA analyses were repeated for SNPs nominally associated with cognition (*p* < 0.05), but generally not associated with education (*p* > 0.05), and then repeated again with the stricter threshold for education (*p* > 0.50). Values for r_g_ of -.55 and -.10 were obtained between schizophrenia and these cognition subsets (Figure 2a).

The second approach involved calculating the heterogeneity p-values for cognition and education, identifying SNPs that have discrepant direction of effects between cognition and education. These SNPs are then binned, ranging from low probability (p < 0.5) to high probability (p < 0.001) for heterogeneous effect sizes between cognition and education (Figure 2b). GNOVA indicated that the greater the discrepancy in effect direction between SNP effects for cognition and education, the stronger the association between cognition and schizophrenia. The same pattern was observed for education and schizophrenia, although to a more modest degree.

### Stage 2: ASSET Meta-Analysis and SNP subsets

Genome-wide cross-phenotype ASSET meta-analysis across 7,306,098 SNPs revealed 300 lead SNPs across 236 independent loci that met the genomewide significance threshold of p <5×10^−8^ for the ASSET 2-tailed test (see Figure 3a and *Supplementary* Tables 1 and 2). There were 1,381,020 SNPs that demonstrated consistent direction of effects between cognition, education and schizophrenia (i.e., lower cognitive ability, lower educational attainment, and increased risk for schizophrenia); these were assigned to the “Concordant” subset, which contained 89 genomewide significant loci harboring 103 independent significant SNPs. By contrast, the “Discordant” subset, which consisted of SNPs with counter-intuitive allelic effects for schizophrenia vis-à-vis education, encompassed 1,891,743 SNPs, with 65 genomewide significant loci comprising 77 independent significant SNPs (Figures 3b and 3c; *Supplementary* Table 1). Significant loci for other ASSET subsets are also detailed in Supplementary Table 2, along with FUMA-derived annotations for potential functional consequences, including CADD scores (*Supplementary* Table 3), eQTL lookups (*Supplementary* Table 4), and prior GWAS lookups (*Supplementary* Table 5).

### Consolidation of Independent Loci to Identify Novel Hits

Next, we wanted to identify which loci from our ASSET results were novel with respect to the three input GWAS. Using RAggr^45^ we extracted SNPs with r^2^ > 0.6 within a window of 250kb of lead SNPs in reported GWAS i.e. 101 loci from the European-ancestry cohorts of the Psychiatric Genomics Consortium GWAS of schizophrenia^33^, 74 loci from the SSGAC educational attainment GWAS^34^ and 40 loci from the COGENT GWAS of cognitive ability^19^. These were merged with the 236 loci from ASSET. As earlier described, independent loci within 250kb were merged, resulting in 280 independent loci being identified across ASSET and the input GWAS. As shown in the resulting Venn diagram (Figure 4), 110 “novel” loci were identified by the ASSET meta-analysis. By contrast, 126 loci overlapped with either schizophrenia, education or schizophrenia, while 44 loci were only significant in the input GWAS but not ASSET.

Very recently, new GWAS have been published for schizophrenia, cognitive ability, and educational attainment, which are larger than the input GWAS used for our ASSET analysis^22,35,36^. This permitted us to perform a lookup of our 110 “novel” ASSET SNPs, thus providing an opportunity to validate ASSET as a tool for novel locus discovery (*Supplementary* Table 6). We also performed lookup in a paper utilizing MTAG to examine intelligence^20^ or several recent papers using pleiotropic approaches to these phenotypes^21,31^. We found that 75% of the loci were in fact reported as significant in the later GWASs with larger sample sizes, while 28 of the 110 loci were novel. The 28 novel loci are reported in Table 1. Further ANNOVAR^43^ annotations are available for novel loci (*Supplementary Table* 7).

### MAGMA Gene-based Analysis: Tissue expression and Competitive Pathway Analysis

MAGMA gene-based analysis was conducted on all ASSET subsets. 772 genes survived Bonferroni correction in the overall ASSET analysis, with 306 genes in the Concordant subset and 304 genes within the Discordant subset (*Supplementary* Table 8). MAGMA gene property analysis revealed significant association (p < 0.000926, Bonferroni-corrected) of gene expression of ASSET SNP subsets across GTExv7 brain tissues (*Supplementary* Figure 3, *Supplementary* Table 9). There were no significant differences between Concordant and Discordant result subsets; both subsets were significantly enriched (positive Beta weights) across all brain compartments.

Because of the significant enrichment in brain tissues, we next performed MAGMA competitive pathway analyses using neurodevelopmental and other brain-related gene sets as curated in a recent publication (Singh et al. 2017); full results are reported in *Supplementary* Table 10. Although there was considerable overlap of pathway enrichment across ASSET categories, several gene sets were uniquely associated with either the Concordant or Discordant result subsets (Table 2). Specifically, the CHD8 pathway, reflecting genes involved in early neurodevelopment, was uniquely associated with the Concordant subset (*p* = 7.11×10^−6^). By contrast, a number of synaptic pathways (e.g. ion channel, synaptic density) and constrained gene sets appeared to be uniquely associated with the Discordant subset. It is notable that when the MHC region is removed from the pathway analysis, the overall pattern of results remained (See *Supplementary* Table 10).

To see if the Concordant/Discordant distinction harbors potential implications for drug targeting (for schizophrenia and/or cognitive enhancement), we performed drug-based and drug family competitive gene set analysis on our MAGMA results. These analyses revealed that the class of drugs associated with voltage-gated calcium channel genes was over-represented amongst the Discordant subset results (Bonferroni-corrected p=0.02), as was Abacavir (nucleoside reverse transcriptase inhibitor; Bonferroni-corrected p=0.00018). While both of these sets showed similar direction of effects with respect to the Concordant subset, no drug-related gene sets attained Bonferroni-corrected significance in the Concordant set of results (Supplementary Table 11).

### S-Predixcan: Brain tissue expression profiles and gene-set enrichment analysis

Transcriptome-wide association analysis (TWAS) was carried out via S-Predixcan to identify top expressed genes within GTEXv7^48^ and CommonMind Consortium^71,74–76^ brain tissue models (Figure 5; *Supplementary* Table 12). The top brain expressed genes unique to the Discordant subsets were *CYP21A1P, CFB*, and *C4A*, along with 177 additional genes significantly expressed in the Discordant, but not Concordant, subsets. On the other hand *ELOV7, NAGA* and 201 other genes were uniquely associated with the Concordant subset. Significant genes identified by S-Predixcan were subjected to gene-set analysis using GENE2FUNC hypergeometric gene set analysis (excluding MHC genes, which were over-represented due to significant linkage disequilibrium, see Methods for more details). The goal of this analysis was to examine if the genes identified in the TWAS overlapped with those found in known biological systems. As shown in Table 3, the results of the TWAS consistently identified genes found in cell adhesion and membrane protein gene sets for the Concordant subset. By contrast, synaptic (specifically, dendritic) pathways, as well as chromosomal repair pathways were consistently identified by the TWAS when examining the Discordant subset.

### Genetic Correlations

A series of psychiatric, personality, structural brain imaging, and metabolic, cardiovascular and anthropometric traits were selected for GNOVA modelling with the ASSET subset results (See Figure 6, *Supplementary* Table 13); multiple testing was adjusted using false discovery rate (FDR). The Concordant subset demonstrated significant (FDR<.05) genetic correlations, in the expected direction, with many forms of psychopathology in addition to schizophrenia (ADHD, bipolar disorder, and MDD, as well as neuroticism and smoking). This subset also demonstrated a significant (FDR<.05) positive genetic correlation (i.e., better cognition/higher education/lower risk for schizophrenia) with larger volumes of several brain regions (including total intracranial volume) as measured by structural MRI, as well as several measures of infant size and adult height. Significant positive associations were also seen with the personality dimensions of openness and conscientiousness, and (surprisingly) self-reported cancer; significant negative associations were seen for total cholesterol and triglycerides, as well as presence of ulcerative colitis/inflammatory bowel disease. Additionally, a negative genetic correlation was observed for the Concordant subset with BMI and measures of cardiovascular disease (i.e., lower cognition/lower education/greater risk for schizophrenia associated with greater BMI and risk for cardiovascular disease).

The Discordant subset was strongly associated with schizophrenia and education, by definition, in a manner demonstrating the paradoxical relationship (higher education, greater risk for schizophrenia (Figure 6). (It is important to note that the light blue bars and dark blue bars in Figure 6 are essentially mirror images of each other and are therefore providing somewhat redundant information; both sets of bars are included to indicate the both sides of this dimension). Interestingly, a similar pattern was observed for bipolar disorder (higher education/greater risk for schizophrenia – greater risk for bipolar disorder). Similar relationships were also observed, at a nominally significant level, for autism spectrum disorder and eating disorders, which were not associated with the concordant subset, as well as MDD. The reverse relationship, however, was observed with ADHD (i.e., higher education/greater risk for schizophrenia – lower risk for ADHD). This pattern was also observed for the smoking, BMI, and cardiovascular disease phenotypes. A counter-intuitive pattern was observed for the relationship between the Discordant subset and neuroticism, which was the opposite of that observed for MDD (despite the fact that MDD and neuroticism are themselves highly correlated).

## Discussion

A consistent finding in recent schizophrenia, cognitive, and educational GWAS has been the involvement of both neurodevelopmental pathways and synaptic processes^19,20,34,78,79^; the present study aimed to at least partially disentangle these mechanisms. In this study, we leveraged the genetic pleiotropy underlying three partially overlapping, complex phenotypes in order to identify homogeneous subsets of SNPs with distinct characteristics. Specifically, we were able to parse a subset of SNPs with alleles that were associated in the expected fashion across our three phenotypes of interest: lower cognitive ability, lower educational attainment, and greater risk for schizophrenia. These “Concordant” SNPs were characterized by their association with genes and pathways relevant to early neurodevelopmental processes. By contrast, SNPs that demonstrated a counterintuitive, discordant pattern of association (higher educational attainment yet greater risk for schizophrenia), were primarily associated with genes/pathways involved in synaptic function of mature neurons.

This distinction was robustly observed across several methods of functional annotation. First, MAGMA competitive gene-set analysis revealed a significant enrichment of CHD8-related genes in the Concordant subset (Table 2). *CHD8*, encoding a chromatin remodeling protein, is a gene that has been robustly associated with autism^80–83^, but to date has only limited or anecdotal evidence for association to schizophrenia^84,85^. Disruption of the homologous gene (*Chd8*) in animal models has demonstrated that the resulting protein plays a key role in very early neurodevelopmental processes, including neuronal proliferation and differentiation^86,87^, as well as cell adhesion and axon guidance^88^. On the other hand, MAGMA competitive gene-set analysis revealed a significant enrichment of discordant genes for functions including synaptic transmission and the postsynaptic density, as well as membrane depolarization and voltage-gated cation channel activity. While these processes have been commonly associated with both schizophrenia^33,35^ and cognitive phenotypes^22–24,90–92^, our study is the first to demonstrate that these synaptic mechanisms operate in a surprising manner: the same synaptic functions that increase risk for schizophrenia also serve to enhance educational attainment.

The linkage of early neurodevelopmental processes to SNPs associated with impaired cognition and increased risk for schizophrenia is consistent with a large literature demonstrating that cognitive deficits are often observed early on in the lifespan of these individuals^13,14,93,94^. At the same time, the Discordant variant subset implicates more mature neuronal regulation, and synaptic pruning mechanisms that are most prominent later in childhood, adolescence, and into adulthood, ostensibly as part of a neuroplasticity mechanism to make more “efficient” connections within the brain^95^. However, the dysregulation of such mechanisms have been shown to be intricately linked to schizophrenia psychopathology^96^. It is important to note that these results are obtained from separate GWASs of two different phenotypes, and do not represent a subset of highly educated patients with schizophrenia. Rather, it is plausible to posit an inverted U relationship, such that efficient synaptic pruning processes are essential mechanisms underlying academic performance, but may be carried too far in disorders such as schizophrenia.

Additionally, transcriptome-wide analysis using S-Predixcan pointed towards the same distinction between Concordant and Discordant genes and pathways. Two of the strongest genes with differential expression in the Concordant subset were *NAGA* (an enzyme cleaving specific moieties from glycoconjugates) and *NDUFAF2* (part of the mitochondrial complex); rare mutations in each of these genes are associated with early and severe neurodevelopmental disorders^97,98^. TWAS of the discordant subset revealed synaptic genes including *C4A*, which plays a key role in synaptic pruning^96^, as well as other transcripts essential to synapse structure and function such as *ARL3, FXR1*, and *CNNM2.* Moreover, pathway analysis of S-Predixcan results (Table 3) demonstrated that the strongest gene set associated with the Concordant subset was cell-cell adhesion via plasma-membrane adhesion molecules (GO:0098742), which encompasses processes such necessary for neural tube closure, cerebral cortex migration, and neuronal-glial interactions. By contrast, the Discordant subset transcriptome was significantly enriched for genes located at dendrites, as well as DNA repair. Recently, the role of DNA repair in modulating neuronal activity-induced gene expression has been shown to be crucial for synaptic plasticity and related processes of learning and memory^99^; impairments in DNA repair have been linked to neurodegeneration^100,101^

ASSET also permitted the identification of novel SNPs for cognition-related phenotypes. Lookups of the full ASSET results revealed that ∼75% of the additional 110 loci, not identified in the input GWAS studies^19,33,34^, were in fact replicated in by an MTAG study examining intelligence^20^ and more recent follow-up GWAS^22,35,36^ with larger samples that were more well powered for variant discovery. This result strongly supports the validity of the ASSET methodology, and demonstrates that the approach indeed improves power for cross-phenotype discovery of new loci as previous discussed by the developers of the method^32^. Notably, several of our novel loci were associated with eQTLs suggesting new potential biological mechanisms for individual variation in cognitive and psychiatric phenotypes. For example, one of the novel loci strongly implicates variation in *PLXNB2*, a gene associated with GABA and glutamate synapses in the hippocampus^102^. Another novel locus shows strong eQTL signal with *NDE1*, a neurodevelopmental gene at the 16p13.11 locus, where copy number variants have been associated with neurodevelopmental disorders^103^.

Our work supports and extends a recent study by Bansal and colleagues^30^, which is the only published report (to our knowledge) that has deeply examined the paradoxical relationship between educational attainment and schizophrenia. Using a proxy-phenotype approach, these investigators identified two novel loci, implicating the *FOXO6* and *SLITRK1* genes, with pleiotropic (i.e., “discordant”) effects across the two phenotypes. Using ASSET, we also uncovered those genes amongst our 110 “novel” loci (one of which was also not identified in any of the updated single-phenotype GWAS, see Table 1). Several other studies^19–21,31^ have employed other statistical approaches to identify pleiotropy and/or overlap across cognitive/educational and schizophrenia GWAS, uncovering a subset of the novel loci identified by ASSET. By utilizing ASSET, we were able to systematically and powerfully identify concordant and discordant pleiotropic loci across the genome, and to then characterize underlying biological mechanisms.

In addition to functional characterization using pathway analyses, we were able to characterize the Concordant and Discordant SNP sets with respect to genetic overlap with other relevant phenotypes. To our knowledge, this is the first study to examine genetic correlations with dimensional sub-sets, rather than global correlations with full GWAS. While the Concordant subset followed the expected patterns of genetic correlation with several forms of psychopathology, as well as brain/head size, results for the Discordant subset were somewhat surprising. For example, we had anticipated that the Discordant subset might be significantly related to personality, as a non-cognitive trait that could promote greater educational attainment. However, correlations with conscientiousness, openness, and neuroticism were stronger for the Concordant as opposed to the Discordant subset.

On the other hand, significant correlations for the Discordant subset were observed with risk for autism, which has previously been shown to demonstrate a counter-intuitive positive genetic correlation with cognition^104^. Given that variants within the Discordant subset tend to index regulation of synaptic function and pruning processes, our results suggest that these mechanisms be investigated with respect to their impact on autism, eating disorders, and bipolar disorder. Moreover, it is noteworthy that autism, despite being a neurodevelopmental disorder, did not demonstrate a significant genetic correlation with the Concordant subset, indicating that it does not share the specific neurodevelopmental pathways implicated in the common variant genetic overlap between schizophrenia risk and impaired cognition. It is also intriguing that bipolar disorder demonstrated a very similar pattern of GNOVA results to schizophrenia, despite prior reports that bipolar disorder is not significantly correlated at the genetic level with general cognitive ability^104,105^. Thus, our approach was able to refine components of neurodevelopment and synaptic function that are shared across cognitive phenotypes, schizophrenia, and bipolar disorder. Further research is needed to identify components of cognition that differentiate schizophrenia and bipolar disorder.

One limitation of this study is only common SNPs (MAF > 0.01) were examined. The genetic architecture of cognitive ability and educational attainment is composed of causal variants in LD with common SNPs (cognitive ability h^2^ = 22.7%, education h^2^ = 15.6%) as well as with causal variants in LD with rare and less common SNPs (cognitive ability h^2^ = 31.3%, education h^2^ = 28.1%), with rarer variants making greater contribution to cognitive differences than more common variants^106^. Rare variants are also known to explain some of the differences in schizophrenia prevalence^52^. However, GNOVA, used to identify genetic correlations across data independent data sets using summary GWAS data, can only capture the contributions made by common genetic effects. Future work aiming to investigate the concordant and discordant effect of rare variants across cognitive ability, schizophrenia, and education is needed^107^. Additionally, the input GWAS for ASSET were of somewhat different sample sizes and power, with the cognitive GWAS demonstrating smaller mean effect sizes compared to schizophrenia and educational attainment; the effects of such differences on ASSET results are not fully understood, although ASSET has been benchmarked as the best available approach to handling non-uniform distribution of effect sizes^40^.

Having demonstrated the utility and validity of the ASSET approach, future studies are planned that can further exploit this method using larger, and more varied, input GWAS. Recent studies have demonstrated that genetic correlations exist across seemingly disparate brain-related phenotypes.^108^ However, such genetic correlations only describe the average genetic effect between pairs of traits. As such, they are not informative as to which variants are associated across traits, nor if a minority of these variants have effects across traits that are the opposite of what would be expected by the direction of the genetic correlation. The application of the ASSET approach to these data sets would help to move beyond the analysis of shared genetic variance, and begin to identify shared genetic variants which, as shown in the current study, may be composed of variants with different combinations of protective and deleterious effects. Future studies, with additional statistical techniques, incorporating rare variants, and novel annotation resources, are needed to further decompose the early neurodevelopmental and adult synaptic pathways highlighted in the present report.

## Supporting information

Main Tables

Main Figures

## Figure Legends

**Figure 1.** Design of the present study. a) input GWAS studies used for ASSET analysis. b) definition of Concordant and Discordant SNP subsets. Concordant SNPs have alleles which demonstrate negative effects on cognitive ability, educational attainment, and schizophrenia risk (i.e., increased schizophrenia risk, reverse-coded for consistency). Discordant SNPs have alleles which demonstrate paradoxical effects on educational attainment and schizophrenia (i.e., higher educational attainment and increased schizophrenia risk, reverse-coded).

**Figure 2.** Genetic correlations with schizophrenia, for SNPs demonstrating heterogeneity of effects between cognitive ability and educational attainment.

**Figure 3.** Manhattan plots for ASSET results. a) All subsets. b) Concordant subset. c) Discordant subset

**Figure 4.** Venn diagram comparing significant ASSET loci to significant loci from input GWASs.

**Figure 5.** Transcriptome-wide association results using S-PrediXcan as applied to Concordant and Discordant subsets.

**Figure 6.** Genetic correlations for Concordant and Discordant subsets with other relevant phenotypes.

## Acknowledgments

This work has been supported by grants from the National Institutes of Health (R01 MH079800 and P50 MH080173 to AKM; R01 MH080912 to DCG; K23 MH077807 to KEB; K01 MH085812 to MCK). Data collection for the TOP cohort was supported by the Research Council of Norway, South-East Norway Health Authority, and KG Jebsen Foundation. The NCNG study was supported by Research Council of Norway Grants 154313/V50 and 177458/V50. The NCNG GWAS was financed by grants from the Bergen Research Foundation, the University of Bergen, the Research Council of Norway (FUGE, Psykisk Helse), Helse Vest RHF and Dr Einar Martens Fund. The Helsinki Birth Cohort Study has been supported by grants from the Academy of Finland, the Finnish Diabetes Research Society, Folkhälsan Research Foundation, Novo Nordisk Foundation, Finska Läkaresällskapet, Signe and Ane Gyllenberg Foundation, University of Helsinki, Ministry of Education, Ahokas Foundation, Emil Aaltonen Foundation. For the LBC1936 cohort, phenotype collection was supported by The Disconnected Mind project. Genotyping was funded by the UK Biotechnology and Biological Sciences Research Council (BBSRC grant No. BB/F019394/1). The work was undertaken by The University of Edinburgh Centre for Cognitive Ageing and Cognitive Epidemiology, part of the cross council Lifelong Health and Wellbeing Initiative, which is funded by the Medical Research Council and the Biotechnology and Biological Sciences Research Council (MR/K026992/1). WDH is supported by a grant from Age UK (Disconnected Mind Project). The CAMH work was supported by the CAMH Foundation and the Canadian Institutes of Health Research. The Duke Cognition Cohort (DCC) acknowledges K. Linney, J.M. McEvoy, P. Hunt, V. Dixon, T. Pennuto, K. Cornett, D. Swilling, L. Phillips, M. Silver, J. Covington, N. Walley, J. Dawson, H. Onabanjo, P. Nicoletti, A. Wagoner, J. Elmore, L. Bevan, J. Hunkin and R. Wilson for recruitment and testing of subjects. DCC also acknowledges the Ellison Medical Foundation New Scholar award AG-NS-0441-08 for partial funding of this study as well as the National Institute of Mental Health of the National Institutes of Health under award number K01MH098126. The UCLA Consortium for Neuropsychiatric Phenomics (CNP) study acknowledges the following sources of funding from the NIH: Grants UL1DE019580 and PL1MH083271 (RMB), RL1MH083269 (TDC), RL1DA024853 (EL) and PL1NS062410. The ASPIS study was supported by National Institute of Mental Health research grants R01MH085018 and R01MH092515 to Dr. Dimitrios Avramopoulos. Support for the Duke Neurogenetics Study was provided the National Institutes of Health (R01 DA033369 and R01 AG049789 to ARH) and by a National Science Foundation Graduate Research Fellowship to MAS. Recruitment, genotyping and analysis of the TCD healthy control samples were supported by Science Foundation Ireland (grants 12/IP/1670, 12/IP/1359 and 08/IN.1/B1916). Data access for several cohorts used in this study was provided by the National Center for Biotechnology Information (NCBI) database of Genotypes and Phenotypes (dbGaP). dbGaP accession numbers for these cohorts were:

Cardiovascular Health Study (CHS): phs000287.v4.p1, phs000377.v5.p1, and phs000226.v3.p1

Framingham Heart Study (FHS): phs000007.v23.p8 and phs000342.v11.p8

Multi-Site Collaborative Study for Genotype-Phenotype Associations in Alzheimer’s Disease (GENADA): phs000219.v1.p1

Long Life Family Study (LLFS): phs000397.v1.p1

Genetics of Late Onset Alzheimer’s Disease Study (LOAD): phs000168.v1.p1

Minnesota Center for Twin and Family Research (MCTFR): phs000620.v1.p1

Philadelphia Neurodevelopmental Cohort (PNC): phs000607.v1.p1

The acknowledgment statements for these cohorts are found below:

Framingham Heart Study: The Framingham Heart Study is conducted and supported by the National Heart, Lung, and Blood Institute (NHLBI) in collaboration with Boston University (Contract No. N01-HC-25195 and HHSN268201500001I). This manuscript was not prepared in collaboration with investigators of the Framingham Heart Study and does not necessarily reflect the opinions or views of the Framingham Heart Study, Boston University, or NHLBI. Funding for SHARe Affymetrix genotyping was provided by NHLBI Contract N02-HL-64278. SHARe Illumina genotyping was provided under an agreement between Illumina and Boston University.

Cardiovascular Health Study: This research was supported by contracts HHSN268201200036C, HHSN268200800007C, N01-HC-85079, N01-HC-85080, N01-HC-85081, N01-HC-85082, N01-HC-85083, N01-HC-85084, N01-HC-85085, N01-HC-85086, N01-HC-35129, N01 HC-15103, N01 HC-55222, N01-HC-75150, N01-HC-45133, and N01-HC-85239; grant numbers U01 HL080295 and U01 HL130014 from the National Heart, Lung, and Blood Institute, and R01 AG-023629 from the National Institute on Aging, with additional contribution from the National Institute of Neurological Disorders and Stroke. A full list of principal CHS investigators and institutions can be found at https://chs-nhlbi.org/pi. This manuscript was not prepared in collaboration with CHS investigators and does not necessarily reflect the opinions or views of CHS, or the NHLBI. Support for the genotyping through the CARe Study was provided by NHLBI Contract N01-HC-65226. Support for the Cardiovascular Health Study Whole Genome Study was provided by NHLBI grant HL087652. Additional support for infrastructure was provided by HL105756 and additional genotyping among the African-American cohort was supported in part by HL085251, DNA handling and genotyping at Cedars-Sinai Medical Center was supported in part by National Center for Research Resources grant UL1RR033176, now at the National Center for Advancing Translational Technologies CTSI grant UL1TR000124; in addition to the National Institute of Diabetes and Digestive and Kidney Diseases grant DK063491 to the Southern California Diabetes Endocrinology Research Center.

Multi-Site Collaborative Study for Genotype-Phenotype Associations in Alzheimer’s Disease: The genotypic and associated phenotypic data used in the study were provided by the GlaxoSmithKline, R&D Limited. Details on data acquisition have been published previously in: Li H, Wetten S, Li L, St Jean PL, Upmanyu R, Surh L, Hosford D, Barnes MR, Briley JD, Borrie M, Coletta N, Delisle R, Dhalla D, Ehm MG, Feldman HH, Fornazzari L, Gauthier S, Goodgame N, Guzman D, Hammond S, Hollingworth P, Hsiung GY, Johnson J, Kelly DD, Keren R, Kertesz A, King KS, Lovestone S, Loy-English I, Matthews PM, Owen MJ, Plumpton M, Pryse-Phillips W, Prinjha RK, Richardson JC, Saunders A, Slater AJ, St George-Hyslop PH, Stinnett SW, Swartz JE, Taylor RL, Wherrett J, Williams J, Yarnall DP, Gibson RA, Irizarry MC, Middleton LT, Roses AD. Candidate single-nucleotide polymorphisms from a genomewide association study of Alzheimer disease. Arch Neurol., Jan;65(1):45-53, 2008 (PMID: 17998437).

Filippini N, Rao A, Wetten S, Gibson RA, Borrie M, Guzman D, Kertesz A, Loy-English I, Williams J, Nichols T, Whitcher B, Matthews PM. Anatomically-distinct genetic associations of APOE epsilon4 allele load with regional cortical atrophy in Alzheimer’s disease. Neuroimage, Feb 1;44(3):724-8, 2009. (PMID: 19013250).

Genetics of Late Onset Alzheimer’s Disease Study: Funding support for the “Genetic Consortium for Late Onset Alzheimer’s Disease” was provided through the Division of Neuroscience, NIA. The Genetic Consortium for Late Onset Alzheimer’s Disease includes a genome-wide association study funded as part of the Division of Neuroscience, NIA. Assistance with phenotype harmonization and genotype cleaning, as well as with general study coordination, was provided by Genetic Consortium for Late Onset Alzheimer’s Disease. A list of contributing investigators is available at https://www.ncbi.nlm.nih.gov/projects/gap/cgi-bin/study.cgi?study_id=phs000168.v1.p1 Long Life Family Study: Funding support for the Long Life Family Study was provided by the Division of Geriatrics and Clinical Gerontology, National Institute on Aging. The Long Life Family Study includes GWAS analyses for factors that contribute to long and healthy life. Assistance with phenotype harmonization and genotype cleaning as well as with general study coordination, was provided by the Division of Geriatrics and Clinical Gerontology, National Institute on Aging. Support for the collection of datasets and samples were provided by Multicenter Cooperative Agreement support by the Division of Geriatrics and Clinical Gerontology, National Institute on Aging (UO1AG023746; UO1023755; UO1023749; UO1023744; UO1023712). Funding support for the genotyping which was performed at the Johns Hopkins University Center for Inherited Disease Research was provided by the National Institute on Aging, National Institutes of Health.

Minnesota Center for Twin and Family Research: This project was led by William G. Iacono, PhD. And Matthew K. McGue, PhD (Co-Principal Investigators) at the University of Minnesota, Minneapolis, MN, USA. Co-investigators from the same institution included: Irene J. Elkins, Margaret A. Keyes, Lisa N. Legrand, Stephen M. Malone, William S. Oetting, Michael B. Miller, and Saonli Basu. Funding support for this project was provided through NIDA (U01 DA 024417). Other support for sample ascertainment and data collection came from several grants: R37 DA 05147, R01 AA 09367, R01 AA 11886, R01 DA 13240, R01 MH 66140.

Philadelphia Neurodevelopmental Cohort: Support for the collection of the data sets was provided by grant RC2MH089983 awarded to Raquel Gur, MD, and RC2MH089924 awarded to Hakon Hakonarson, MD, PhD. All subjects were recruited through the Center for Applied Genomics at The Children’s Hospital in Philadelphia.

## Available Software

ASSET https://dceg.cancer.gov/tools/analysis/asset

LDSC https://github.com/bulik/ldsc

FUMA http://fuma.ctglab.nl/

MAGMA https://ctg.cncr.nl/software/magma

S-Predixcan https://github.com/hakyimlab/MetaXcan

VEP http://grch37.ensembl.org/Homo_sapiens/Tools/VEP

