## Supplementary material for "Pleiotropic Meta-Analysis of Cognition, Education, and Schizophrenia Differentiates Roles of Early Neurodevelopmental and Adult Synaptic Pathways": Main Tables

| **Table 1**  **Novel Loci Identified by ASSET** | | | |  |  | |  | |  | |  | |  | |
| --- | --- | --- | --- | --- | --- | --- | --- | --- | --- | --- | --- | --- | --- | --- |
| Lead SNPs | chr | pos_start | pos_end | | | gwasP | | beta | | se | | nearestGene | | func |
| rs13010104 | 2 | 208331258 | 208369715 | | | 9.30E-09 | | -0.0148 | | 0.0026 | | ENSG00000223725 | | ncRNA_intronic |
| rs207338 | 4 | 19053350 | 19070123 | | | 4.99E-08 | | -0.0108 | | 0.0020 | | ENSG00000248238 | | intergenic |
| rs6844280 | 4 | 31371970 | 31387144 | | | 3.53E-08 | | 0.0113 | | 0.0021 | | ENSG00000251434 | | intergenic |
| rs71615297 | 4 | 179013479 | 179210420 | | | 2.17E-08 | | -0.0115 | | 0.0021 | | RNU1-45P | | intergenic |
| rs73260443 | 5 | 113404038 | 113464825 | | | 6.30E-09 | | -0.0147 | | 0.0025 | | ENSG00000251628 | | intergenic |
| rs9372208 | 6 | 109507533 | 109648130 | | | 2.96E-09 | | -0.0120 | | 0.0020 | | ENSG00000233908 | | intergenic |
| rs9371912 | 6 | 155907483 | 156009420 | | | 1.06E-10 | | -0.0168 | | 0.0026 | | RNU7-152P | | intergenic |
| rs1870571 | 8 | 80129136 | 80296429 | | | 4.69E-09 | | 0.0124 | | 0.0021 | | ENSG00000253659 | | intergenic |
| rs11786685 | 8 | 81220185 | 81316928 | | | 2.79E-08 | | -0.0116 | | 0.0021 | | ENSG00000253237 | | intergenic |
| rs3890699 | 8 | 110164392 | 110344596 | | | 7.31E-09 | | 0.0122 | | 0.0021 | | NUDCD1 | | intergenic |
| rs11166628 | 8 | 137022220 | 137091947 | | | 3.27E-08 | | 0.0113 | | 0.0020 | | ENSG00000253248 | | ncRNA_intronic |
| rs10993909 | 9 | 136924744 | 136942560 | | | 3.13E-08 | | 0.0116 | | 0.0021 | | BRD3 | | intronic |
| rs10994707 | 10 | 62919886 | 63096664 | | | 1.14E-09 | | 0.0202 | | 0.0033 | | TMEM26 | | intergenic |
| rs72945305 | 11 | 81164016 | 81210528 | | | 3.34E-08 | | -0.0128 | | 0.0023 | | ENSG00000254747 | | intergenic |
| rs556587 | 11 | 92317365 | 92554716 | | | 1.57E-08 | | -0.0163 | | 0.0029 | | FAT3 | | intronic |
| rs75261250 | 11 | 124276497 | 124303201 | | | 2.16E-08 | | -0.0188 | | 0.0034 | | OR8X1P | | intergenic |
| rs708212 | 12 | 31517960 | 31769501 | | | 7.79E-09 | | 0.0133 | | 0.0023 | | DENND5B | | intronic |
| rs3741434 | 12 | 53605344 | 53605344 | | | 1.20E-10 | | 0.0191 | | 0.0030 | | RARG | | UTR3 |
| rs7321596 | 13 | 44390857 | 44514022 | | | 3.16E-08 | | -0.0109 | | 0.0020 | | LACC1 | | intergenic |
| rs11617058 | 13 | 85176690 | 85305386 | | | 2.95E-11 | | 0.0183 | | 0.0028 | | LINC00333 | | intergenic |
| rs67652508 | 14 | 55487496 | 55567991 | | | 2.88E-08 | | -0.0140 | | 0.0025 | | MAPK1IP1L | | intergenic |
| rs11130 | 16 | 15687755 | 15837246 | | | 4.24E-08 | | 0.0110 | | 0.0020 | | NDE1 | | UTR3 |
| rs7214058 | 17 | 9968014 | 9995284 | | | 6.32E-09 | | 0.0118 | | 0.0020 | | GAS7 | | intronic |
| rs28584904 | 17 | 68984046 | 69007006 | | | 2.36E-08 | | 0.0144 | | 0.0026 | | ENSG00000271101 | | intergenic |
| rs56791590 | 18 | 26259012 | 26496051 | | | 7.59E-09 | | 0.0117 | | 0.0020 | | ENSG00000265994 | | intergenic |
| rs12462428 | 19 | 16665215 | 16738369 | | | 2.20E-08 | | 0.0143 | | 0.0026 | | ENSG00000141979:MED26:CTC-429P9.4 | | intronic:intronic:intronic |
| rs5767976 | 22 | 48133458 | 48183889 | | | 8.14E-10 | | -0.0125 | | 0.0020 | | RP11-191L9.4 | | ncRNA_intronic |
| rs68178377 | 22 | 50742346 | 50771464 | | | 4.28E-08 | | 0.0122 | | 0.0022 | | DENND6B | | exonic |

| **Table 2** |  |  |  | |  |  |  |
| --- | --- | --- | --- | --- | --- | --- | --- |
| MAGMA Significant Gene Sets for Concordant and Discordant SNP Subsets | | |  | |  |  |  |
| MAGMA Gene Sets | NGENES | P | | Pbon | BETA | BETA_STD | SE |
| *CONCORDANT* | | | | | | | |
| **cotney_2015_hNSC_Chd8_prom** | **8186** | **4.05E-09** | | **7.11E-06** | **0.115** | **0.0571** | **0.0199** |
| cotney_2015_hNSC+human+mouse_Chd8_prom | 1902 | 2.81E-05 | | 0.049458 | 0.123 | 0.0378 | 0.0304 |
| sugathan_2014_chd8_binding | 5314 | 1.69E-05 | | 0.029744 | 0.0874 | 0.04 | 0.0211 |
| *DISCORDANT* | | | | | | | |
| **constrained_genes_0_10** | **1705** | **3.41E-07** | | **0.0006** | **0.131** | **0.0383** | **0.0263** |
| **constrained_genes_pLI_90** | **2972** | **3.33E-07** | | **0.000585** | **0.104** | **0.0387** | **0.021** |
| **darnell_2011_fmrp_targets** | **765** | **1.41E-05** | | **0.024693** | **0.162** | **0.0327** | **0.0387** |
| ddg2p_dominant_mis_all_brain | 137 | 4.42E-06 | | 0.007768 | 0.411 | 0.0357 | 0.0925 |
| **g2cdb_bayes_collins_mouse_psd_consensus** | **918** | **8.89E-06** | | **0.015615** | **0.152** | **0.0333** | **0.0353** |
| **g2cdb_bayes_collins_mouse_psd_full** | **1442** | **1.26E-06** | | **0.002214** | **0.134** | **0.0363** | **0.0284** |
| **g2cdb_human_psd** | **1001** | **1.40E-06** | | **0.002461** | **0.159** | **0.0363** | **0.0339** |
| **g2cdb_human_psp** | **1039** | **2.04E-06** | | **0.003579** | **0.154** | **0.0358** | **0.0334** |
| GOBP:membrane_depolarization | 105 | 2.17E-05 | | 0.038129 | 0.439 | 0.0334 | 0.107 |
| GOBP:synaptic_transmission | 549 | 6.58E-06 | | 0.011554 | 0.205 | 0.0353 | 0.0471 |
| **GOMF:voltage-gated_cation_channel_activity** | **144** | **6.43E-07** | | **0.00113** | **0.45** | **0.0401** | **0.093** |
| **sanders_2015_asd_fdr10** | **64** | **4.67E-06** | | **0.008207** | **0.622** | **0.037** | **0.14** |
| **sanders_2015_asd_lof_genes** | **559** | **3.52E-06** | | **0.006178** | **0.203** | **0.0352** | **0.0451** |
| **Results remaining significant after removal of MHC region variants are highlighted in Bold** | | | | | | | |

| **Table 3** |  |  |  |  |  |
| --- | --- | --- | --- | --- | --- |
| GENE2FUNC Pathway analysis of GO genesets with MHC filtered | |  |  |  |  |
| Category | GeneSet | N_genes | N_overlap | p | Adj. P |
| *CONCORDANT* | |  |  |  |  |
| GO_bp | GO_CELL_CELL_ADHESION_VIA_PLASMA_MEMBRANE_ADHESION_MOLECULES | 202 | 10 | 6.89E-09 | 3.1E-05 |
| GO_bp | GO_RESPONSE_TO_XENOBIOTIC_STIMULUS | 105 | 7 | 6.22E-08 | 2.8E-04 |
| GO_bp | GO_HOMOPHILIC_CELL_ADHESION_VIA_PLASMA_MEMBRANE_ADHESION_MOLECULES | 151 | 8 | 7.74E-08 | 3.4E-04 |
| GO_bp | GO_CELL_CELL_ADHESION | 604 | 14 | 4.46E-07 | 2.0E-03 |
| GO_mf | GO_OXIDOREDUCTASE_ACTIVITY_ACTING_ON_NAD_P_H_QUINONE_OR_SIMILAR_COMPOUND_AS_ACCEPTOR | 52 | 4 | 6.39E-06 | 5.8E-03 |
| GO_cc | GO_MEMBRANE_MICRODOMAIN | 286 | 8 | 1.53E-05 | 8.9E-03 |
| GO_cc | GO_MEMBRANE_PROTEIN_COMPLEX | 1018 | 16 | 1.58E-05 | 9.2E-03 |
| GO_bp | GO_PURINE_RIBONUCLEOSIDE_BISPHOSPHATE_METABOLIC_PROCESS | 20 | 3 | 2.77E-06 | 1.2E-02 |
| GO_cc | GO_MITOCHONDRION | 1633 | 21 | 2.47E-05 | 1.4E-02 |
| GO_bp | GO_BIOLOGICAL_ADHESION | 1027 | 17 | 4.56E-06 | 2.0E-02 |
| GO_bp | GO_MACROMOLECULAR_COMPLEX_ASSEMBLY | 1388 | 20 | 6.87E-06 | 3.0E-02 |
| GO_bp | GO_MITOCHONDRIAL_RESPIRATORY_CHAIN_COMPLEX_I_BIOGENESIS | 56 | 4 | 9.25E-06 | 4.1E-02 |
| *DISCORDANT* | |  |  |  |  |
| GO_cc | GO_DENDRITIC_SHAFT | 37 | 3 | 1.28E-05 | 7.4E-03 |
| GO_bp | GO_DOUBLE_STRAND_BREAK_REPAIR | 165 | 6 | 3.89E-06 | 1.7E-02 |
| GO_cc | GO_NUCLEAR_CHROMOSOME | 522 | 9 | 3.84E-05 | 2.2E-02 |
| GO_mf | GO_PROTEIN_DOMAIN_SPECIFIC_BINDING | 620 | 10 | 3.14E-05 | 2.8E-02 |
| GO_bp | GO_NON_RECOMBINATIONAL_REPAIR | 70 | 4 | 7.97E-06 | 3.5E-02 |
| GO_cc | GO_DENDRITE | 451 | 8 | 6.94E-05 | 4.0E-02 |
