## Supplementary Figures for "Pleiotropic Meta-Analysis of Cognition, Education, and Schizophrenia Differentiates Roles of Early Neurodevelopmental and Adult Synaptic Pathways"

### Slide 1
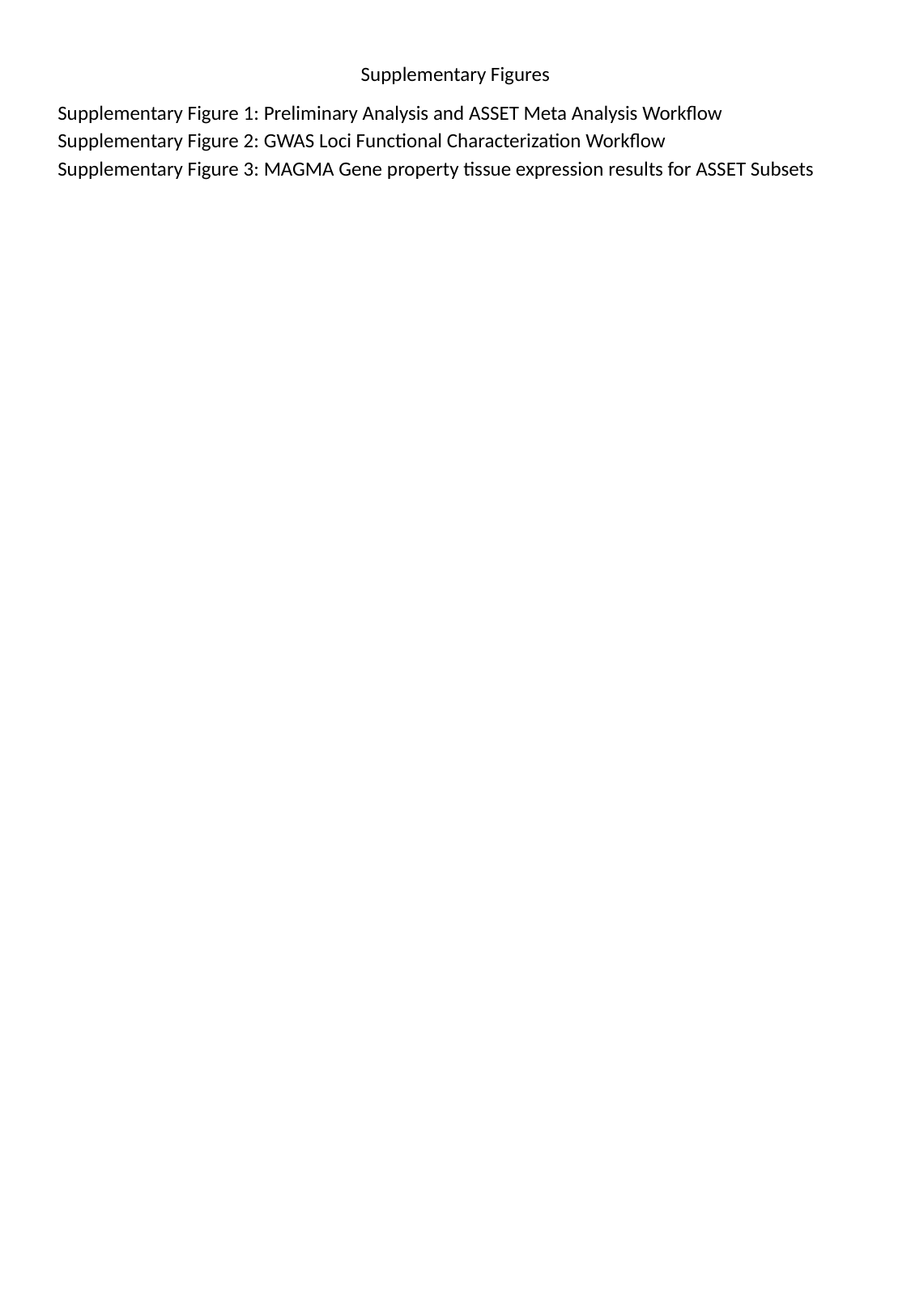

#### Supplementary Figures
Supplementary Figure 1: Preliminary Analysis and ASSET Meta Analysis Workflow
Supplementary Figure 2: GWAS Loci Functional Characterization Workflow
Supplementary Figure 3: MAGMA Gene property tissue expression results for ASSET Subsets

### Slide 2
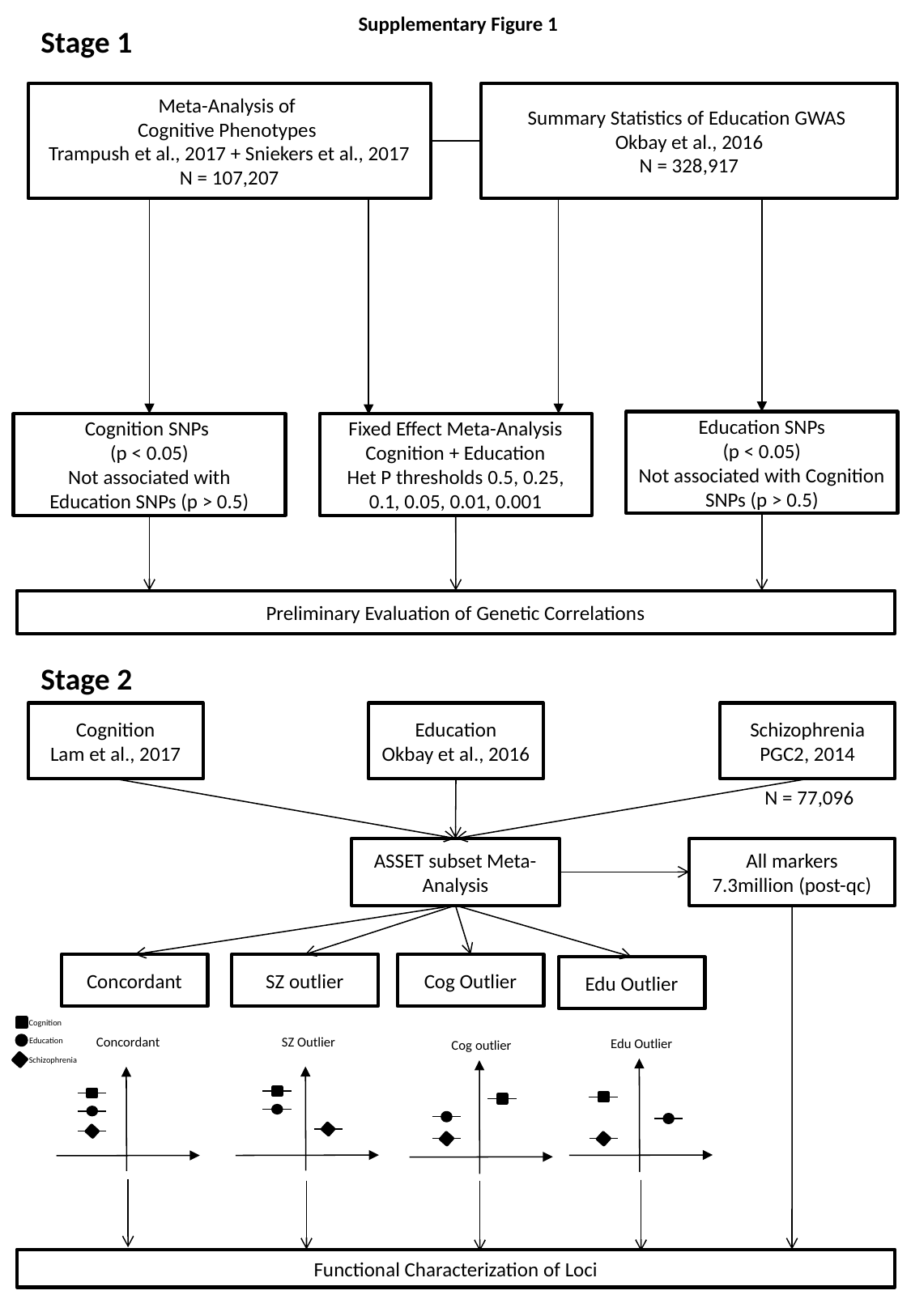

Supplementary Figure 1
Stage 1
Meta-Analysis of
Cognitive Phenotypes
Trampush et al., 2017 + Sniekers et al., 2017
N = 107,207
Summary Statistics of Education GWAS
Okbay et al., 2016
N = 328,917
Education SNPs
(p < 0.05)
Not associated with Cognition SNPs (p > 0.5)
Cognition SNPs
(p < 0.05)
Not associated with Education SNPs (p > 0.5)
Fixed Effect Meta-Analysis
Cognition + Education
Het P thresholds 0.5, 0.25, 0.1, 0.05, 0.01, 0.001
Preliminary Evaluation of Genetic Correlations
Stage 2
Education
Okbay et al., 2016
Schizophrenia
PGC2, 2014
Cognition
Lam et al., 2017
N = 77,096
ASSET subset Meta-Analysis
All markers
7.3million (post-qc)
Concordant
SZ outlier
Cog Outlier
Edu Outlier
Cognition
Education
Schizophrenia
SZ Outlier
Concordant
Edu Outlier
Cog outlier
Functional Characterization of Loci

### Slide 3
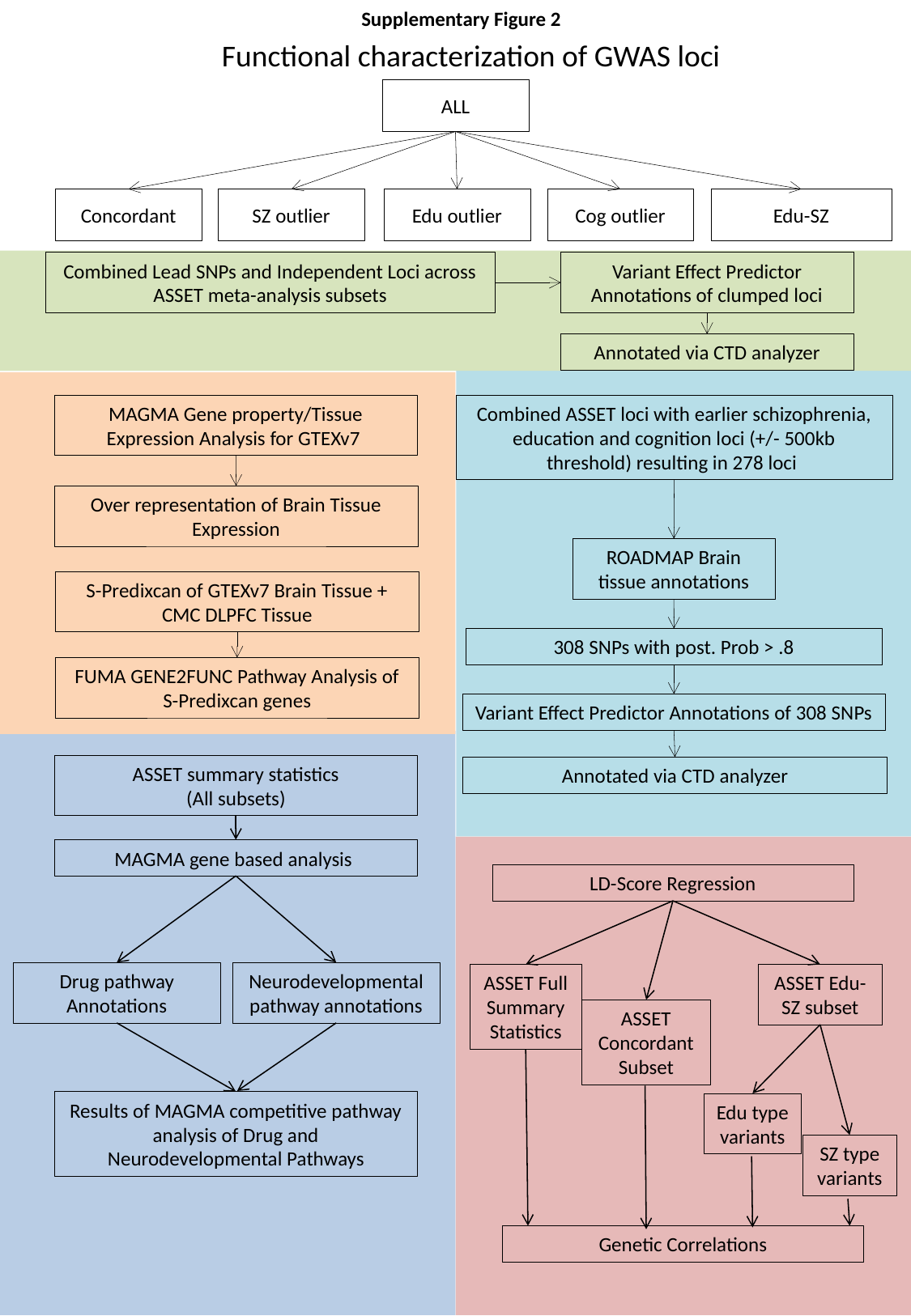

Supplementary Figure 2
Functional characterization of GWAS loci
ALL
Edu-SZ
Cog outlier
Concordant
SZ outlier
Edu outlier
Variant Effect Predictor Annotations of clumped loci
Combined Lead SNPs and Independent Loci across ASSET meta-analysis subsets
Annotated via CTD analyzer
MAGMA Gene property/Tissue Expression Analysis for GTEXv7
Combined ASSET loci with earlier schizophrenia, education and cognition loci (+/- 500kb threshold) resulting in 278 loci
Over representation of Brain Tissue Expression
ROADMAP Brain tissue annotations
S-Predixcan of GTEXv7 Brain Tissue + CMC DLPFC Tissue
308 SNPs with post. Prob > .8
FUMA GENE2FUNC Pathway Analysis of S-Predixcan genes
Variant Effect Predictor Annotations of 308 SNPs
ASSET summary statistics
(All subsets)
Annotated via CTD analyzer
MAGMA gene based analysis
LD-Score Regression
Drug pathway Annotations
Neurodevelopmental pathway annotations
ASSET Edu-SZ subset
ASSET Full Summary Statistics
ASSET Concordant Subset
Results of MAGMA competitive pathway analysis of Drug and Neurodevelopmental Pathways
Edu type variants
SZ type variants
Genetic Correlations

### Slide 4
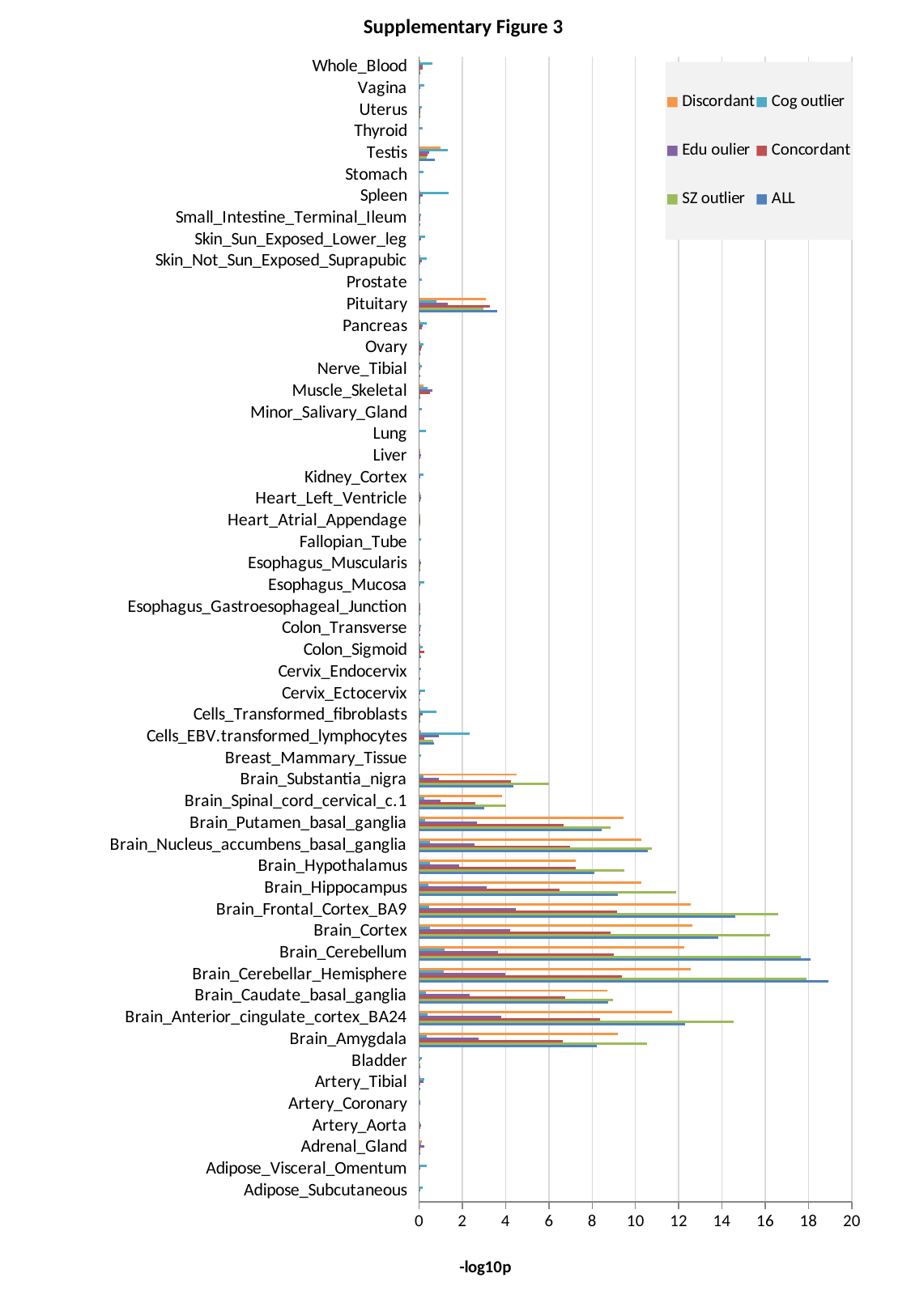

Supplementary Figure 3
###### Chart
| Category | ALL | SZ outlier | Concordant | Edu oulier | Cog outlier | Discordant |
|---|---|---|---|---|---|---|
| Adipose_Subcutaneous | 2.605845067551099e-05 | 0.0 | 1.302902989351207e-05 | 0.009536825119402358 | 0.16771467876513296 | 8.68597649812824e-06 |
| Adipose_Visceral_Omentum | 4.342966533881614e-06 | 0.0 | 0.0 | 0.027117083623571705 | 0.3503341909079744 | 4.342966533881614e-06 |
| Adrenal_Gland | 0.032742274391581554 | 0.02238612507310454 | 0.02815845225928285 | 0.25553489822107667 | 0.10829532376081731 | 0.13367130150604065 |
| Artery_Aorta | 0.0016273063255449322 | 0.00010424318533945632 | 0.002307882058273408 | 0.08707152728457095 | 0.045825052735054335 | 0.0019761862928163324 |
| Artery_Coronary | 0.000408428804615366 | 5.211846499883996e-05 | 6.08054839496669e-05 | 0.023943624409711123 | 0.05128391640249485 | 8.252379157010987e-05 |
| Artery_Tibial | 0.013031310841759016 | 0.001143699111843679 | 0.00468955094531247 | 0.20386990105093847 | 0.23223580550971637 | 0.005300203182952815 |
| Bladder | 0.007539271165210578 | 0.0035321097382827633 | 0.00015637416252358162 | 0.01300446027561836 | 0.13373038758296008 | 0.0002128564502228404 |
| Brain_Amygdala | 8.227084478935293 | 10.517841304588723 | 6.631675918020042 | 2.7493116963542787 | 0.3473186152364258 | 9.20600717793199 |
| Brain_Anterior_cingulate_cortex_BA24 | 12.292327648985252 | 14.546330084934773 | 8.36286064927366 | 3.804155675252718 | 0.3742863721887986 | 11.68235445677884 |
| Brain_Caudate_basal_ganglia | 8.735323817851176 | 8.973385032065325 | 6.769270155422319 | 2.345496175432953 | 0.31927479377584544 | 8.700492701299513 |
| Brain_Cerebellar_Hemisphere | 18.924270600259103 | 17.89445575425343 | 9.359439065595364 | 3.9922095625540215 | 1.1254775959277268 | 12.545109427188763 |
| Brain_Cerebellum | 18.090599103634883 | 17.629931239349947 | 8.988260438611682 | 3.6409997726253134 | 1.1563499514003528 | 12.248921148853828 |
| Brain_Cortex | 13.82637674230192 | 16.233773106425833 | 8.867484165087362 | 4.217369739782391 | 0.5044695378982692 | 12.630914079178492 |
| Brain_Frontal_Cortex_BA9 | 14.622675532263445 | 16.590607969380848 | 9.152439678944564 | 4.470775034372668 | 0.47023609595988225 | 12.560114919182698 |
| Brain_Hippocampus | 9.19099276294457 | 11.880513164435364 | 6.482223670726044 | 3.108892031585053 | 0.44557141527794514 | 10.265792430268544 |
| Brain_Hypothalamus | 8.09820396480079 | 9.49068992579071 | 7.230637446096243 | 1.850565333656831 | 0.4959643296075355 | 7.230674378725626 |
| Brain_Nucleus_accumbens_basal_ganglia | 10.571476459579236 | 10.768224676024905 | 6.9819655972954715 | 2.5543179731473207 | 0.49031370382877126 | 10.27471740715931 |
| Brain_Putamen_basal_ganglia | 8.450959934916531 | 8.843390281834301 | 6.666210261668803 | 2.6611651577089033 | 0.2784861579159776 | 9.445631999009912 |
| Brain_Spinal_cord_cervical_c.1 | 3.0139990384858297 | 4.02886801193393 | 2.596742174212889 | 0.9808837095529273 | 0.24808224451919772 | 3.839051519135303 |
| Brain_Substantia_nigra | 4.3477052992747955 | 6.0062464316351285 | 4.258651397524105 | 0.9070338624056354 | 0.22082213369274534 | 4.510632225295266 |
| Breast_Mammary_Tissue | 4.342966533881614e-06 | 0.0 | 0.0 | 0.0007954869946869879 | 0.08989877078624882 | 0.0 |
| Cells_EBV.transformed_lymphocytes | 0.6705004242371572 | 0.6424844113849032 | 0.2576755782475977 | 0.917969018732988 | 2.327855942981009 | 0.08274663297632398 |
| Cells_Transformed_fibroblasts | 0.01775138705376744 | 0.005806068917770457 | 0.05712917893329934 | 0.15681084046785654 | 0.8093882021863951 | 0.026165837488117443 |
| Cervix_Ectocervix | 0.0028627291427582844 | 0.0007867852679058821 | 9.989921968843116e-05 | 0.009727759904131782 | 0.28435593306204415 | 0.0006084382809090492 |
| Cervix_Endocervix | 0.002351545103793844 | 0.0005692987434380615 | 0.0001998214241273805 | 0.0019412856814944395 | 0.07288125155632043 | 0.00011727534299772331 |
| Colon_Sigmoid | 0.07492523635314526 | 0.03461012978487766 | 0.2549408308025879 | 0.046182341793003324 | 0.17376965698285576 | 0.00683721859568228 |
| Colon_Transverse | 0.00877392430750515 | 0.0016709009975787279 | 0.02391609065626598 | 0.002853985897430567 | 0.08283071302729147 | 0.0003041125891607762 |
| Esophagus_Gastroesophageal_Junction | 0.02041520799833598 | 0.012266719491167767 | 0.04682523628010643 | 0.06860315858461388 | 0.028320667596255324 | 0.001439899084905462 |
| Esophagus_Mucosa | 0.00016071869309628963 | 0.000673678682910036 | 0.00029976662389581986 | 0.043087565090845825 | 0.23763658285492253 | 0.0008477010170804813 |
| Esophagus_Muscularis | 0.02808431685197118 | 0.015886926556722542 | 0.03916310437653804 | 0.07801716116222393 | 0.022994717211356895 | 0.0018976638610393999 |
| Fallopian_Tube | 0.00023023709362944382 | 0.00021720154586423186 | 1.302902989351207e-05 | 0.0007737330046352441 | 0.10342969393805991 | 8.68597649812824e-06 |
| Heart_Atrial_Appendage | 0.0009521478982837452 | 0.0023602782396974493 | 0.014272722870992849 | 0.02990428741117791 | 0.06297887181875171 | 0.007769119755951898 |
| Heart_Left_Ventricle | 0.0009608529390117113 | 0.0017275806142092618 | 0.0543098925568723 | 0.0976000033229123 | 0.08354078730644052 | 0.01940130284334256 |
| Kidney_Cortex | 6.08054839496669e-05 | 9.989921968843116e-05 | 8.68597649812824e-06 | 0.014627423095618814 | 0.19911116198355763 | 0.001326622298790593 |
| Liver | 0.00013465216155356954 | 0.0007171777296113858 | 0.03314520149792774 | 0.1021146078085461 | 0.008260165452788326 | 0.022157551430759585 |
| Lung | 0.0 | 0.0 | 0.0 | 0.0015183387868471403 | 0.3293831135996745 | 0.0 |
| Minor_Salivary_Gland | 4.342966533881614e-06 | 0.0 | 7.383633819013811e-05 | 0.0011916001191059416 | 0.13871656079367548 | 0.0 |
| Muscle_Skeletal | 0.05422131526064368 | 0.0214360484191101 | 0.5051916725915258 | 0.621456752838356 | 0.3922984104189397 | 0.19008356976982263 |
| Nerve_Tibial | 0.005199100705396131 | 0.00048233462017804456 | 0.0015401301074387359 | 0.022276394711152253 | 0.14020145251943436 | 0.0039482579989949815 |
| Ovary | 0.032915580934904785 | 0.01941946865003162 | 0.08819276782883324 | 0.11102343956139162 | 0.19960601403185962 | 0.016793327642295734 |
| Pancreas | 0.0006780283915259828 | 0.0003649606697487103 | 0.13794618019817945 | 0.16950756675091147 | 0.352294363341144 | 0.005128782426078245 |
| Pituitary | 3.607092104706606 | 2.9615388038214365 | 3.2683226123291456 | 1.3112065607144623 | 0.811718520477333 | 3.076279451823367 |
| Prostate | 0.000621485577291545 | 0.0007563305971195446 | 0.0007258780618011368 | 0.0008433496087722727 | 0.11780002346872621 | 0.00023458236316769576 |
| Skin_Not_Sun_Exposed_Suprapubic | 0.0010130868478760507 | 0.001143699111843679 | 0.003917580830575808 | 0.11101783160088018 | 0.35866472727359155 | 0.0023209805110077654 |
| Skin_Sun_Exposed_Lower_leg | 0.0005997403010787693 | 0.0004649438849982298 | 0.0018976638610393999 | 0.08415185396112425 | 0.2864422598640074 | 0.0011785356840623789 |
| Small_Intestine_Terminal_Ileum | 0.005942533611427499 | 0.0009390906643418537 | 0.016107715696685315 | 0.026728945236014812 | 0.07854243462021145 | 0.0007998379234603362 |
| Spleen | 0.014138102439906299 | 0.023750924783278917 | 0.01099093002192709 | 0.166546885268916 | 1.359518563029578 | 0.024315501033689665 |
| Stomach | 0.00023023709362944382 | 3.040167780437153e-05 | 8.252379157010987e-05 | 0.0002215466849785818 | 0.190649066222121 | 0.0 |
| Testis | 0.7117955032484772 | 0.36698627662845584 | 0.40201976479937973 | 0.4491237471571455 | 1.3242124960917208 | 0.9899970258729404 |
| Thyroid | 1.302902989351207e-05 | 8.68597649812824e-06 | 1.302902989351207e-05 | 0.0003606140955401972 | 0.15042090330665023 | 5.64619527538583e-05 |
| Uterus | 0.019801126220023495 | 0.0048783660448882896 | 0.020929885609056953 | 0.012378417874516247 | 0.1144861251469772 | 0.001832239344577682 |
| Vagina | 0.00033888187700458465 | 0.0004519012906030772 | 1.302902989351207e-05 | 0.005775260243128895 | 0.23611548847089872 | 0.0001085871944407351 |
| Whole_Blood | 0.04790531233197986 | 0.06170047991711636 | 0.1462288036875807 | 0.17389927004434344 | 0.6198430484918404 | 0.053184337387099184 |
