## Supplementary material for "Pleiotropic Meta-Analysis of Cognition, Education, and Schizophrenia Differentiates Roles of Early Neurodevelopmental and Adult Synaptic Pathways": Main Figures

### Slide 1
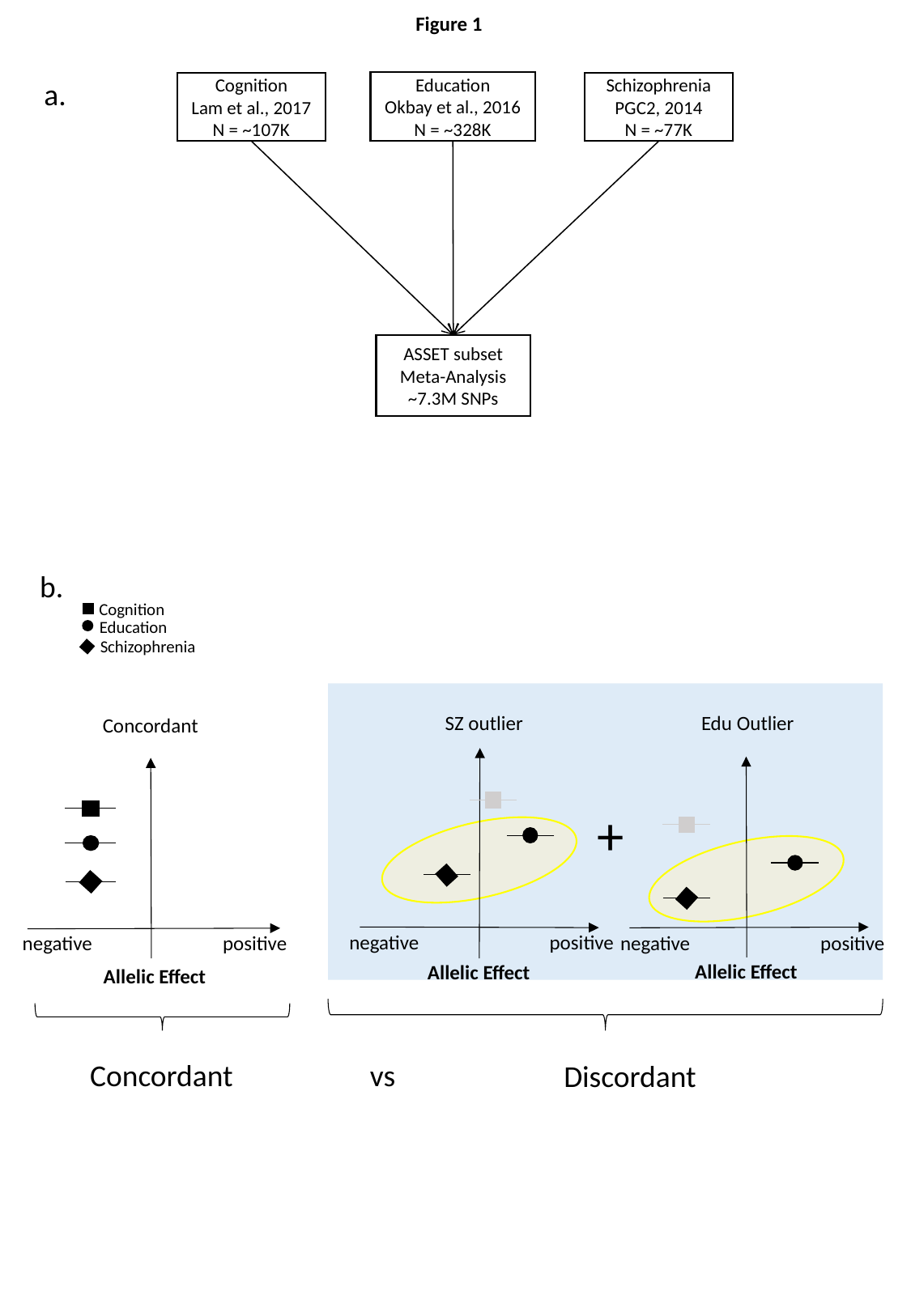

Figure 1
a.
Education
Okbay et al., 2016
N = ~328K
Cognition
Lam et al., 2017
N = ~107K
Schizophrenia
PGC2, 2014
N = ~77K
ASSET subset Meta-Analysis
~7.3M SNPs
b.
Cognition
Education
Schizophrenia
SZ outlier
Edu Outlier
Concordant
+
negative positive
negative positive
negative positive
Allelic Effect
Allelic Effect
Allelic Effect
Concordant
vs
Discordant

### Slide 2
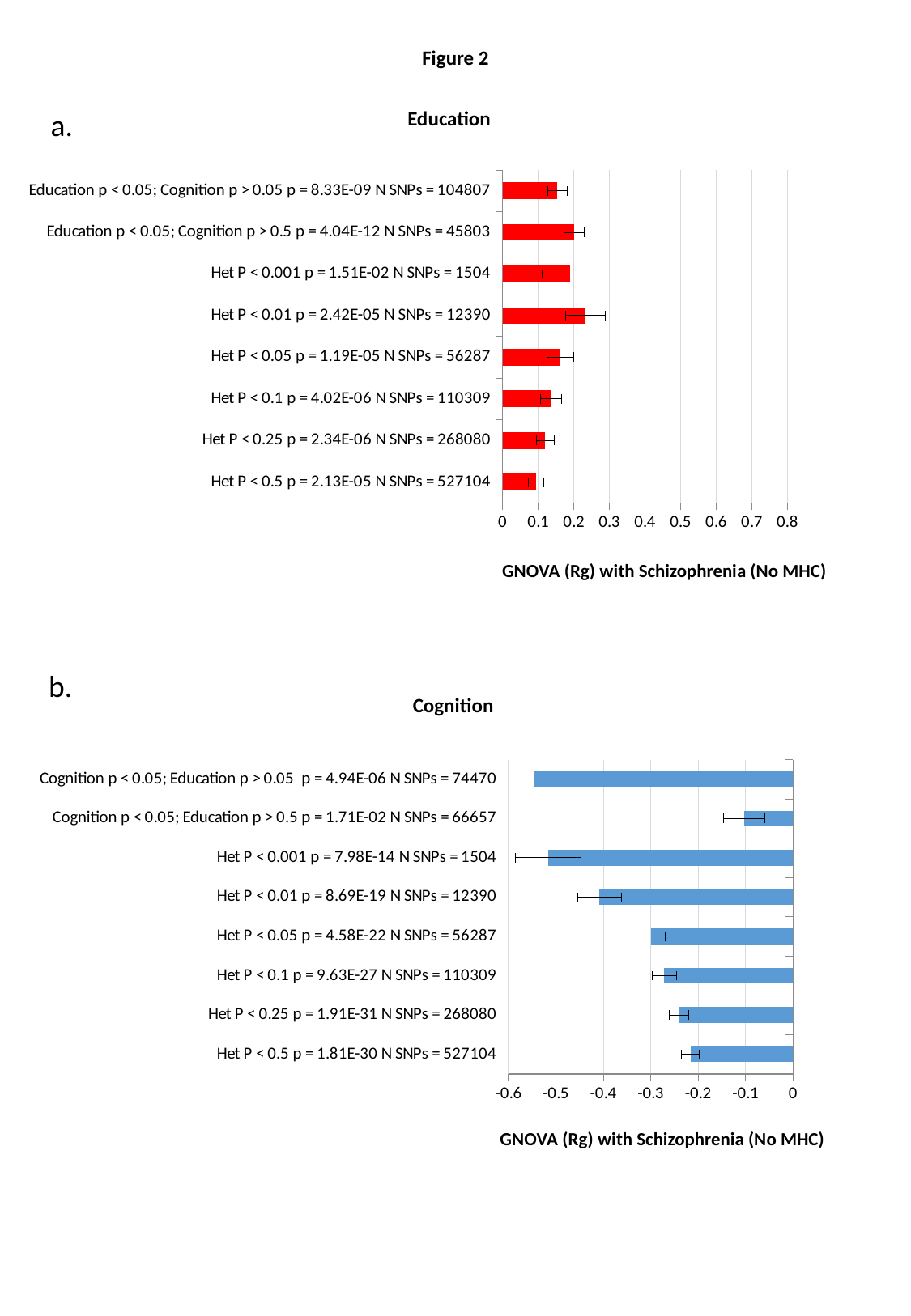

Figure 2
a.
Education
#### Chart
| Category | |
|---|---|
| Het P < 0.5 p = 2.13E-05 N SNPs = 527104 | 0.0942067383350087 |
| Het P < 0.25 p = 2.34E-06 N SNPs = 268080 | 0.12001298294635 |
| Het P < 0.1 p = 4.02E-06 N SNPs = 110309 | 0.136481377031269 |
| Het P < 0.05 p = 1.19E-05 N SNPs = 56287 | 0.16256750793062 |
| Het P < 0.01 p = 2.42E-05 N SNPs = 12390 | 0.233205742939024 |
| Het P < 0.001 p = 1.51E-02 N SNPs = 1504 | 0.189772964321774 |
| Education p < 0.05; Cognition p > 0.5 p = 4.04E-12 N SNPs = 45803 | 0.201249882967099 |
| Education p < 0.05; Cognition p > 0.05 p = 8.33E-09 N SNPs = 104807 | 0.153766335844068 |GNOVA (Rg) with Schizophrenia (No MHC)
b.
Cognition
#### Chart
| Category | |
|---|---|
| Het P < 0.5 p = 1.81E-30 N SNPs = 527104 | -0.216356008444962 |
| Het P < 0.25 p = 1.91E-31 N SNPs = 268080 | -0.240629554645643 |
| Het P < 0.1 p = 9.63E-27 N SNPs = 110309 | -0.271511853134331 |
| Het P < 0.05 p = 4.58E-22 N SNPs = 56287 | -0.300006050218254 |
| Het P < 0.01 p = 8.69E-19 N SNPs = 12390 | -0.408551285657899 |
| Het P < 0.001 p = 7.98E-14 N SNPs = 1504 | -0.516467961507888 |
| Cognition p < 0.05; Education p > 0.5 p = 1.71E-02 N SNPs = 66657 | -0.102959359135993 |
| Cognition p < 0.05; Education p > 0.05 p = 4.94E-06 N SNPs = 74470 | -0.547812497173609 |GNOVA (Rg) with Schizophrenia (No MHC)

### Slide 3
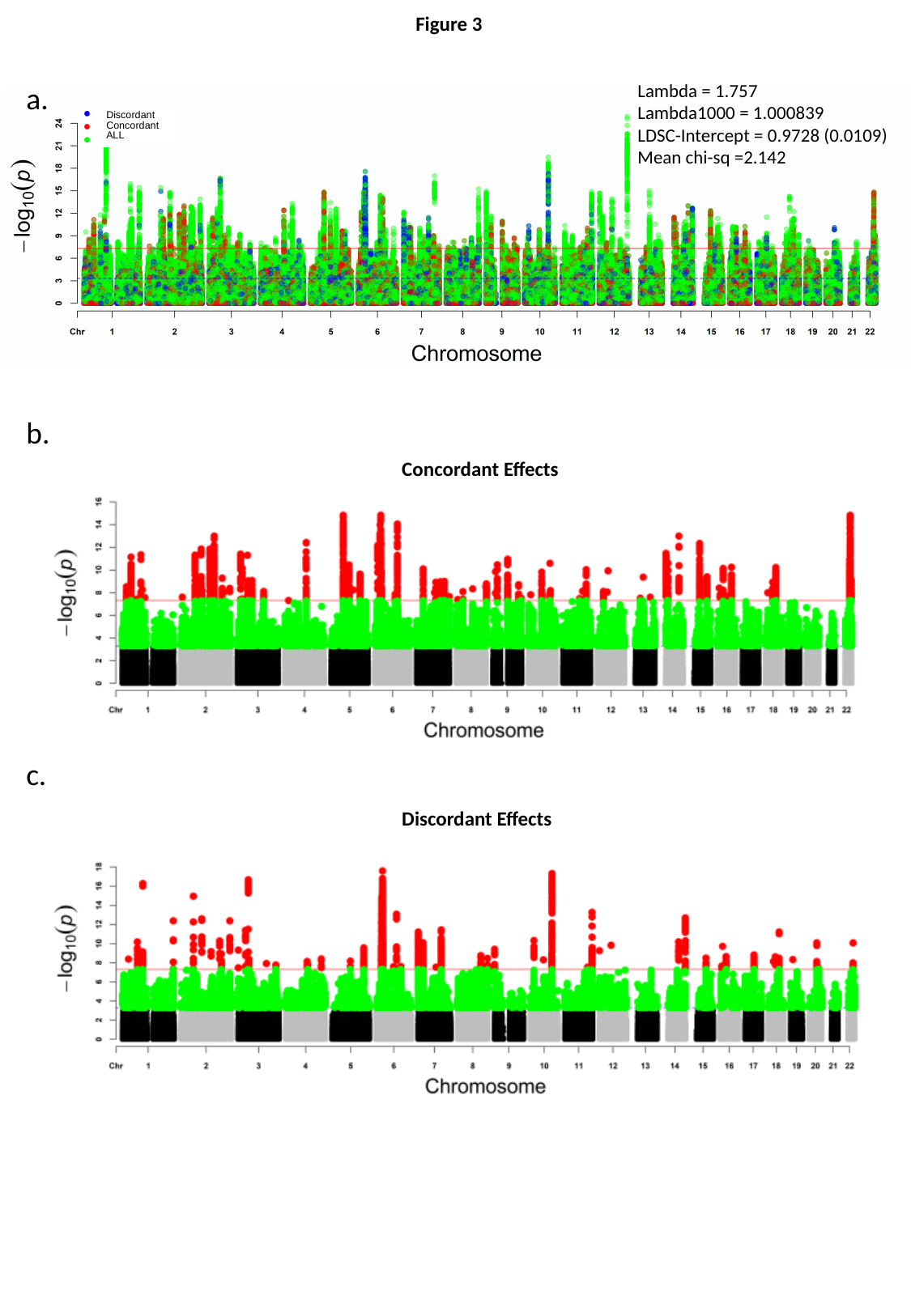

Figure 3
a.
Lambda = 1.757
Lambda1000 = 1.000839
LDSC-Intercept = 0.9728 (0.0109)
Mean chi-sq =2.142
Discordant
Concordant
ALL
b.
Concordant Effects
c.
Discordant Effects

### Slide 4
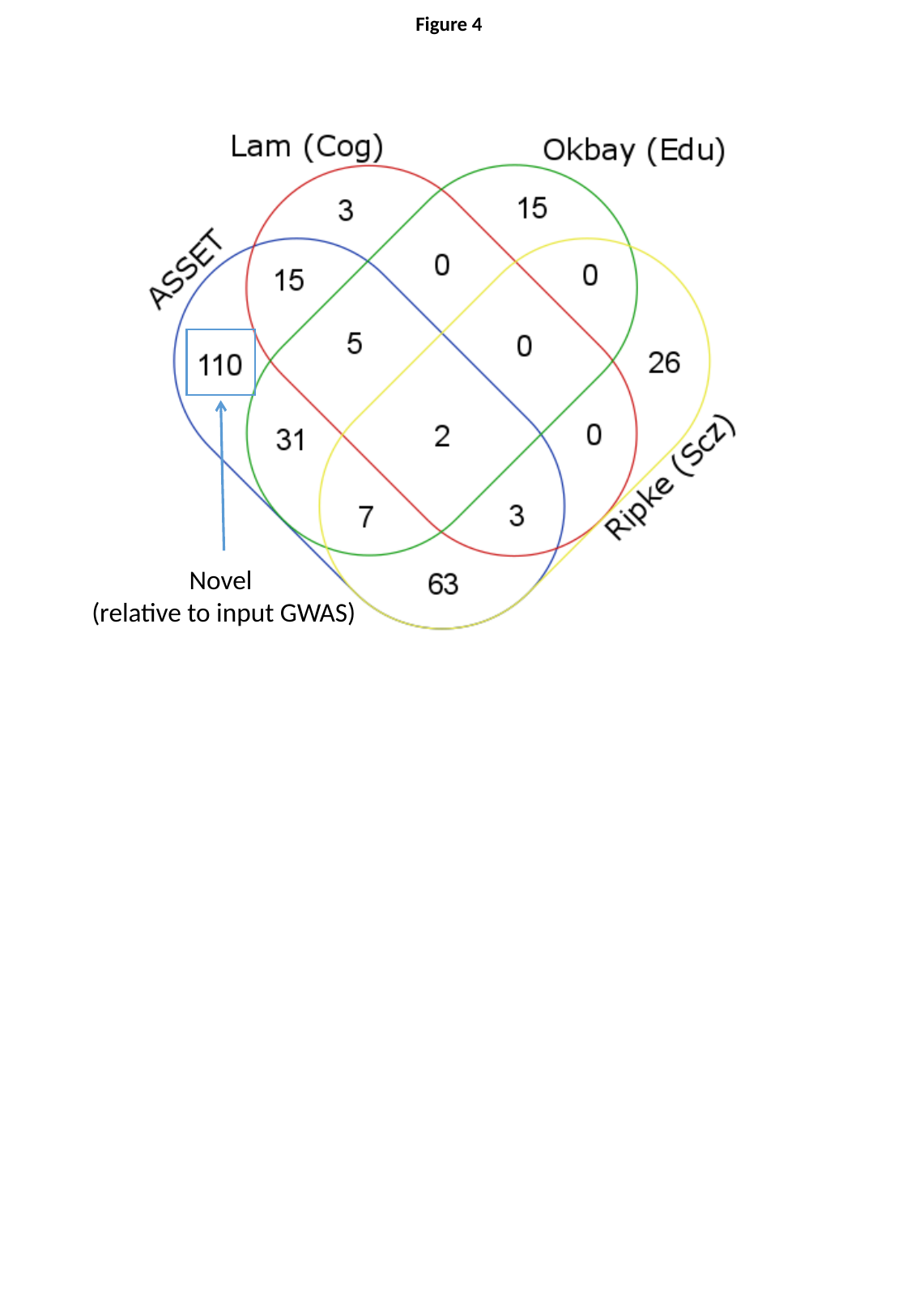

Figure 4
Novel
(relative to input GWAS)

### Slide 5
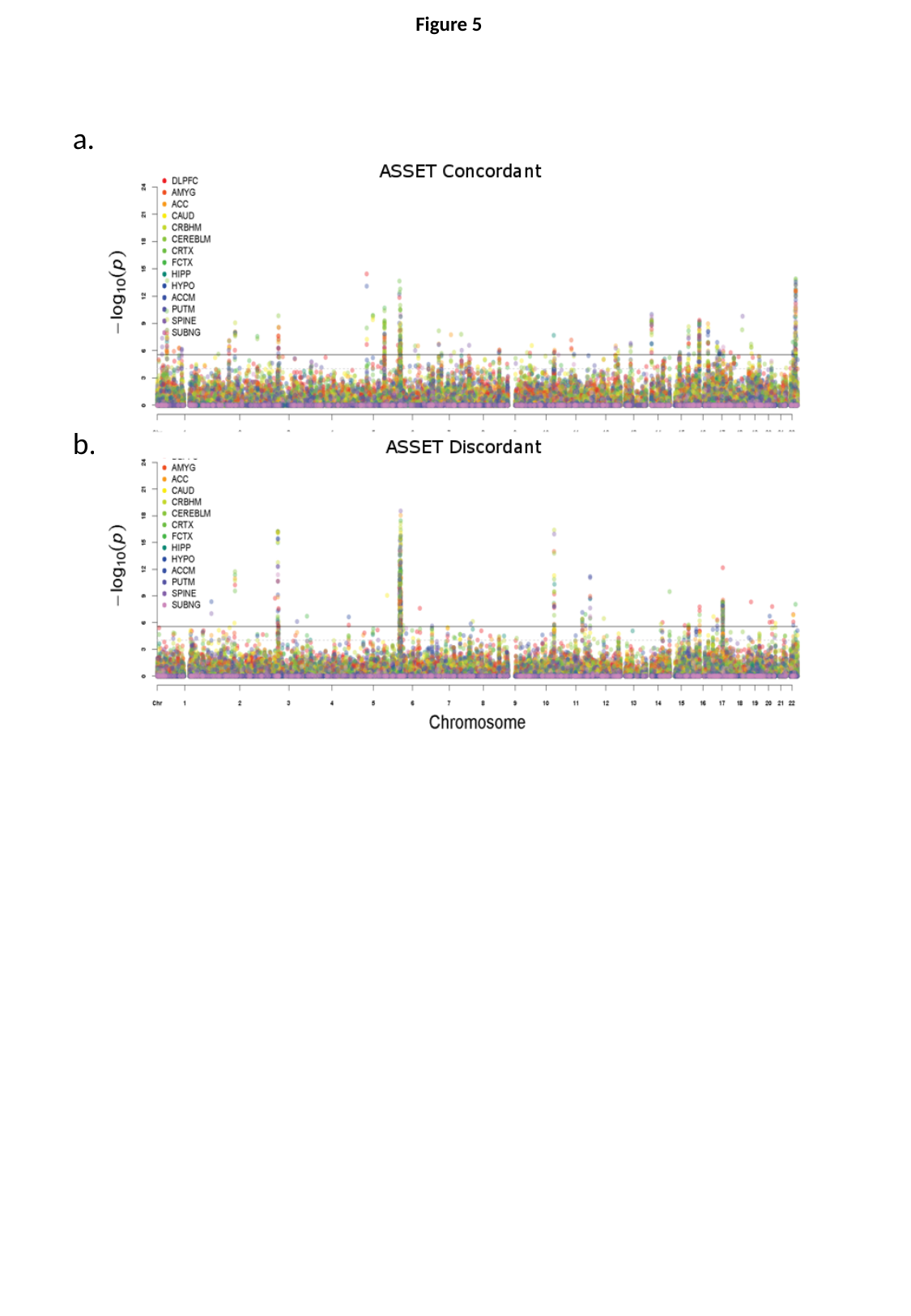

Figure 5
a.
b.

### Slide 6
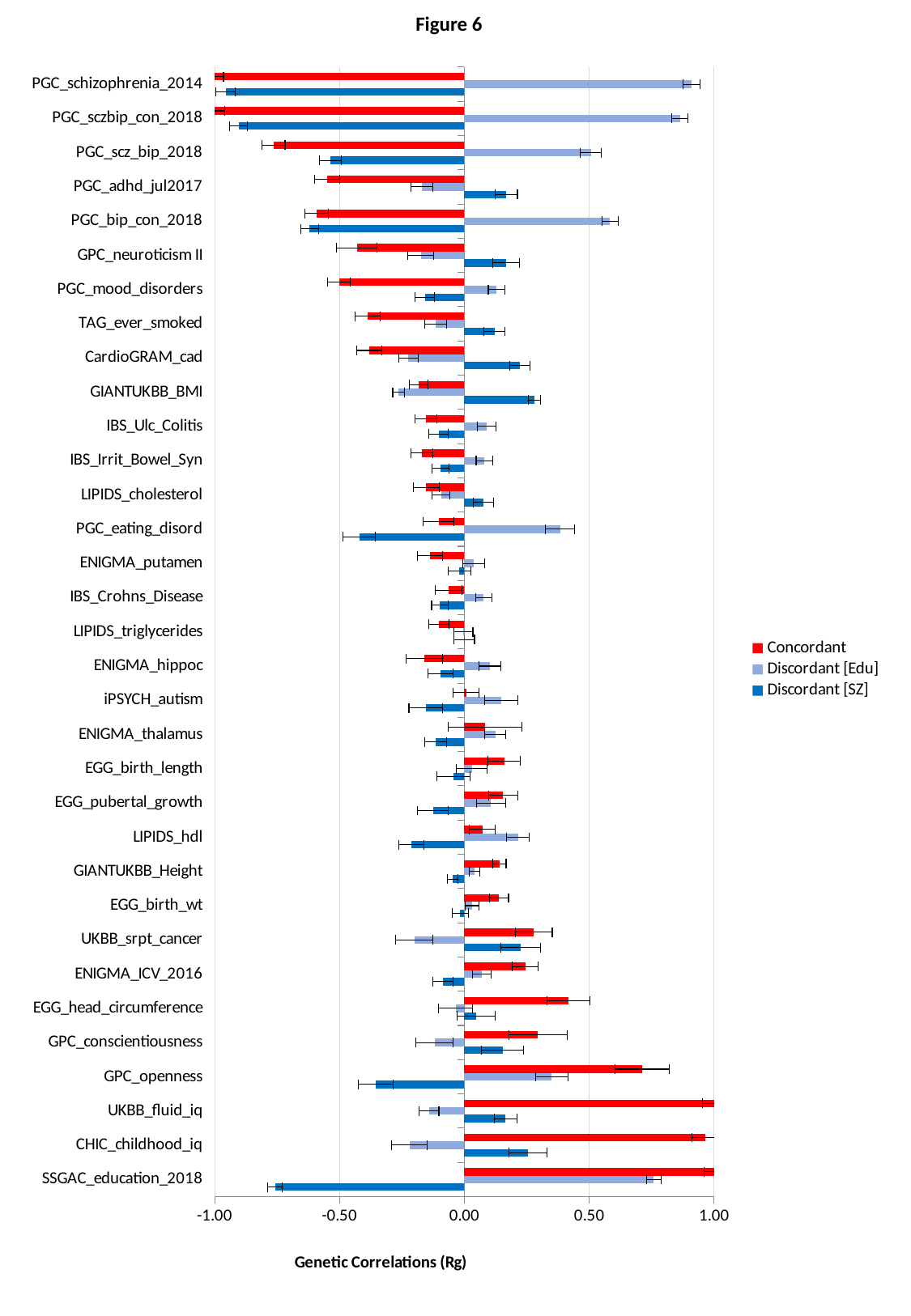

Figure 6
#### Chart
| Category | | | |
|---|---|---|---|
| SSGAC_education_2018 | -0.758207053415972 | 0.758134296973998 | 1.0 |
| CHIC_childhood_iq | 0.255773571313279 | -0.219569310598881 | 0.965175736043676 |
| UKBB_fluid_iq | 0.164478821588133 | -0.141542372930131 | 1.0 |
| GPC_openness | -0.354299313390569 | 0.350313135258636 | 0.711576263191564 |
| GPC_conscientiousness | 0.153577126133295 | -0.11914162518143 | 0.295160959765004 |
| EGG_head_circumference | 0.047264718524609 | -0.0346439021088341 | 0.416428074546789 |
| ENIGMA_ICV_2016 | -0.0860571094893423 | 0.0687283894091306 | 0.244271164499076 |
| UKBB_srpt_cancer | 0.225456766781006 | -0.199537140989246 | 0.278736859492755 |
| EGG_birth_wt | -0.0167023671424053 | 0.0309891021882941 | 0.138495933817254 |
| GIANTUKBB_Height | -0.0454791468877338 | 0.040337601279995 | 0.139902934211976 |
| LIPIDS_hdl | -0.212431460179685 | 0.214684284367435 | 0.0717481725799651 |
| EGG_pubertal_growth | -0.125410361975246 | 0.106567844850655 | 0.154943805133508 |
| EGG_birth_length | -0.0438660426945213 | 0.0295610510178367 | 0.159396604849502 |
| ENIGMA_thalamus | -0.115816575807959 | 0.12428897089037 | 0.0826645760201855 |
| iPSYCH_autism | -0.154566460181614 | 0.148253900359096 | 0.00726085653315264 |
| ENIGMA_hippoc | -0.0959204844198978 | 0.102220151602472 | -0.160122499423646 |
| LIPIDS_triglycerides | -0.000751285816856309 | -0.00405342185295016 | -0.103038001599476 |
| IBS_Crohns_Disease | -0.0976650263455853 | 0.0778802981620474 | -0.0621083834841616 |
| ENIGMA_putamen | -0.0197780971750744 | 0.0377993237784416 | -0.138645415276463 |
| PGC_eating_disord | -0.420638852549544 | 0.383568427754124 | -0.103167129156414 |
| LIPIDS_cholesterol | 0.0767245988679 | -0.0934379288796355 | -0.15247555492287 |
| IBS_Irrit_Bowel_Syn | -0.0963331774902829 | 0.0799511425755546 | -0.169526498807467 |
| IBS_Ulc_Colitis | -0.102833129690544 | 0.089064515880187 | -0.154488999578027 |
| GIANTUKBB_BMI | 0.280816827074827 | -0.263386926458732 | -0.184174319339286 |
| CardioGRAM_cad | 0.222370227590296 | -0.223730715050605 | -0.380736462408765 |
| TAG_ever_smoked | 0.120815036165387 | -0.115255495896869 | -0.387063785138387 |
| PGC_mood_disorders | -0.158420718952709 | 0.129234569974204 | -0.50224471889483 |
| GPC_neuroticism II | 0.167285415705522 | -0.17449184691031 | -0.430869050286945 |
| PGC_bip_con_2018 | -0.619576544778289 | 0.583480214946598 | -0.59195364065951 |
| PGC_adhd_jul2017 | 0.167350693548384 | -0.170098874873887 | -0.549802455140118 |
| PGC_scz_bip_2018 | -0.537472606883676 | 0.506108642916341 | -0.764842004058131 |
| PGC_sczbip_con_2018 | -0.904143718119274 | 0.863318513430911 | -1.0 |
| PGC_schizophrenia_2014 | -0.955932860029373 | 0.909060672234539 | -1.0 |
